# Programmable pattern formation in cellular systems with local signaling

**DOI:** 10.1101/2021.03.17.435764

**Authors:** Tiago Ramalho, Stephan Kremser, Hao Wu, Ulrich Gerland

## Abstract

Diverse complex systems, ranging from developing embryos to systems of locally communicating agents, display an apparent capability of “programmable” pattern formation: They reproducibly form a target pattern, but this target can be readily changed. A distinguishing feature of such systems, as compared to simpler physical pattern forming systems, is that their subunits are capable of information processing. Here, we explore schemes for programmable pattern formation within a theoretical framework, in which subunits process discrete local signals to update their internal state according to logical rules. We study systems with different update rules, different topologies, and different control schemes, to assess their ability to perform programmable pattern formation and their susceptibility to errors. Only a small subset of systems permits local organizer cells to dictate any target pattern. These systems follow a common principle, whereby a temporal pattern is transcribed into a spatial pattern, reminiscent of the clock-and-wavefront mechanism underlying vertebrate somitogenesis. An alternative scheme employing several different rules can only form a fraction of patterns but is robust with respect to the timing of organizer cell inputs. Our results establish a basis for the design of synthetic systems, and for more detailed models of programmable pattern formation closer to real systems.

## INTRODUCTION

Programmable pattern formation is impressively exemplified in developmental biology, where relatively minor changes in the *cis*-regulatory regions of genes can reprogram the developmental process to yield dramatic changes in the morphology of the adult organism^1,2^. In these systems, the individual cells have internal states, but do not know the global state of the system. They process local cues according to their genetic program to determine how and when to change their internal state. Local organizers can induce changes in the internal states of other cells^3^, but there is no global agent overseeing the pattern formation process. Similar behavior can also emerge on a higher level, when e.g. groups of robots^4,5^ or humans^6^ coordinate their motion by local communication. These examples motivate the conceptual question: Which general schemes allow the same agents to produce different complex patterns by following rules to coordinate their behavior with their neighbors?

While natural systems consist of subunits that are already very complex, it is interesting to ask for the simplest model systems capable of programmable pattern formation. Such models would provide a conceptual framework for programmable pattern formation, and could reveal design principles, e.g., for synthetic molecular systems. DNA-based molecular systems, in particular, are readily programmable via the sequence-dependent interaction between DNA strands, which has been exploited to design self-assembling dynamic DNA devices^7^, neural network-like molecular computation^8^, coupled regulatory circuits^9^, and schemes for constructing molecular-scale cellular automata^10^. Here, we use a minimal model to study the concept of programmable pattern formation using theoretical and computational tools. While the intention is not to model any particular system, DNA-based implementations of the model are an interesting perspective (see ‘Discussion’).

We consider a system consisting of spatial subunits with fixed locations on a regular grid. Pattern formation requires the subunits to have at least two distinguishable states. We perform most of our analysis with such minimal subunits, but also present a generalization to subunits with more internal states. The essential model assumptions are that subunits communicate only with their immediate neighbors and that they update their internal states at discrete time steps. The dynamics of a subunit is then governed by update ‘rules’ that depend on its state, as well as on the state of its neighbors. This framework of so-called ‘cellular automata’ is sufficiently flexible to describe a broad range of pattern formation processes that do not depend on long-range signaling between cells^11,12^. Furthermore, cellular automata are not solely abstract computational models, but can faithfully describe the dynamics of real systems, also in developmental biology^13^. For our analysis, a useful feature of cellular automata is that the number of possible update rules is finite – each rule is a different scheme for local information processing – and there are no additional model parameters.

Using this modeling framework, we focus on a scenario in which pattern formation is controlled by ‘organizer cells’, inspired by the Spemann-Mangold organizer in developmental biology^3^. The underlying biological concept is inductive signaling, whereby one cell can change the fate of another cell^14^. For instance, in *Caenorhabditis elegans* vulval patterning, the ‘anchor cell’ controls the cell fate pattern of six vulval precursor cells, involving three different cell fates^15^. Within the conceptual models that we consider, organizer cells can emit time-dependent signals into their neighborhood, affecting the pattern formation process of the remaining ‘bulk cells’. Programmability of pattern formation then refers to the ability of the organizer cells to reproducibly steer the system towards different target patterns, using different signaling sequences. We define a given model to be completely programmable, if organizers can direct the system to all different target patterns from any initial pattern. We consider programmability of pattern formation to be a desirable property, since it is reminiscent of the ability of developmental systems to work with only a small number of signaling systems, which are highly homologous between morphologically very different animals^14^.

Our analysis shows how the dynamics of the bulk cells, as specified by their update rule, affects the programmability of pattern formation. For the minimal system with 2-state cells, only ten update rules enable complete programmability. However, the number of such “programmable rules” increases strongly with the number of internal states. Patterning errors, incurred by cells that do not always follow their update rule, can be strongly reduced by an error correction scheme. The robustness against the timing of organizer inputs can be increased at the expense of a reduction in the extent of programmability if organizer cells are also able to induce changes in the update rules of bulk cells.

## RESULTS

### A minimal model for programmable pattern formation controlled by organizer cells

To explore the programmability of global pattern formation from local sites, we combine concepts from control theory with a class of models for pattern formation. Multiple modeling frameworks for pattern formation processes are available, which treat time, space, and patterning state either as discrete or continuum quantities, and differ also with respect to the level of detail of the description^12^. Here, we choose the most coarse-grained level of description, known as cellular automata (CA) models. Within this framework, a system consists of localized subunits referred to as ‘cells’. The patterning state of cell *i* at time *t* is denoted by 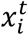, which can only take on a finite number *k* of different values, 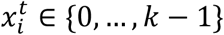. In the simplest case of elementary CA, there are only two different states 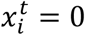 or 1 and the cells are arranged in one dimension (1D). The dynamics of a CA model is governed by local rules specifying how the state of a cell is updated depending on the state of the cell itself and the states of the surrounding cells, see Fig. 1. For elementary CA, only the immediate neighbors of a cell affect its update, via an update rule of the form 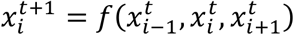. Depending on the update rule, patterning information emerging from a localized source can propagate through the system to affect the global patterning process.

**Figure 1.**
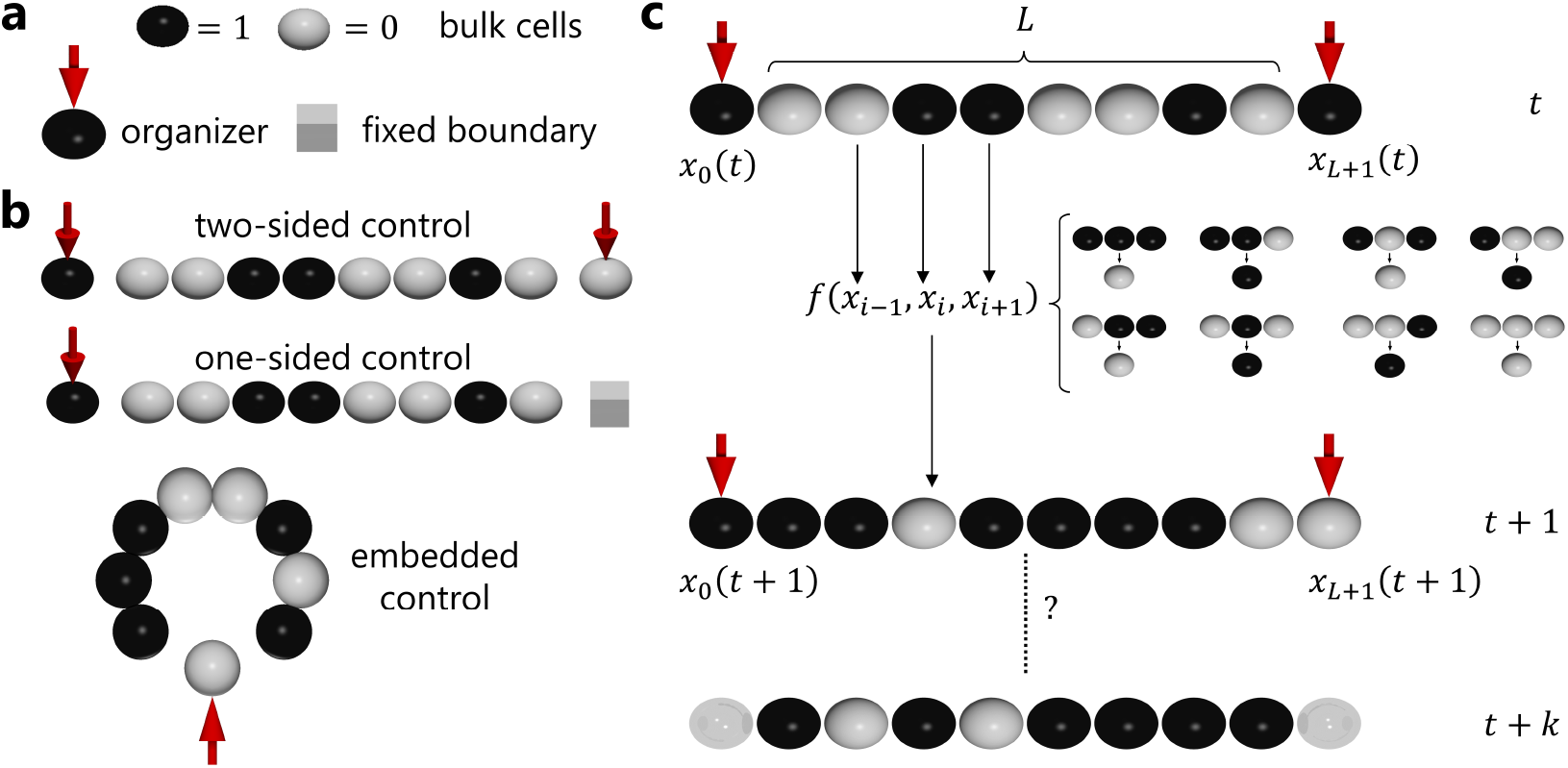
Programmable pattern formation within a cellular automaton (CA) model. **(a)** The model features bulk cells (spheres) and organizer cells (marked by red arrow). The state of cells is represented by their color. Fixed boundary cells (boxes) can also be regarded as organizer cells that never change their state. **(b)** We consider three types of cell arrangement: Linear, with either one or two organizer cells, and circular. **(c)** The patterning dynamics of bulk cells follows a cellular automaton rule. The time-dependent state *x*_*i*_(*t*) of each of the *L* bulk cells is updated according to *x*_*i*_(*t* + 1) = *f*(*x*_*i-1*_(*t*), *x*_*i*_(*t*), *x*_*i-1*_(*t*), with a rule *f* that maps every triple of input states (*x*_*i-1*_(*t*), *x*_*i*_(*t*), *x*_*i-1*_(*t*)) to an output state *x*_*i*_(*t* + 1), as specified by a transition table (small spheres). The rules are enumerated by translating the pattern of output states, ordered by the descending binary equivalent of the input states, into a binary number (here: 01010110_2_, i.e., rule 86). The patterning process is controlled by the local dynamics of the organizer cells, which supply the patterning input. Programmable pattern formation in a cell arrangement with a given update rule refers to the ability of the organizer cell(s) to reproducibly steer the bulk cells to different target patterns, using appropriate sequences of signals (see main text).

A familiar example of global pattern formation from local rules is the performance of large groups of dancers, where the performers produce dynamic patterns by following complex rules for moving in coordination with their immediate neighbors. A global acoustic or optical signal can facilitate the pattern formation process by synchronizing the dynamics. In molecular systems, synchronization can arise from a collectively produced long-range signal, or from a local coupling between oscillators^9^. CA models also typically assume synchronous updates of all cells, and we will follow this convention here. Note that the updates do not need to occur at constant time intervals in real time. Furthermore, the synchronous update assumption is not strictly required, since our model could also be based on an asynchronous CA system with subunits capable of local synchronization via interactions with their neighbors^16^.

Our model systems consist of ‘bulk cells’ and ‘organizer cells’ (Fig. 1a). Bulk cells simply follow their update rules, taking input signals from their neighbor(s), regardless of whether these are other bulk cells or organizer cells. Organizer cells do not take inputs but exert control on the pattern formation process by changing their internal state according to a specified protocol, which depends on the target pattern. We do not specify the origin of this protocol - it could be the result from an internal developmental program or could be the result of external manipulation. Since both natural and engineered patterning systems come in a variety of topologies, and the topology may affect the pattern formation process and its control, we consider two different topologies, linear and circular (Fig. 1b). In a circular topology, a single organizer cell is embedded in a ring of *L* bulk cells, while a linear array of bulk cells can be controlled by an organizer cell at one or both ends, or at any position within the array. We collectively denote the time-dependent patterning state of all bulk cells as *X*^*t*^, and the time-dependent state of the organizer cells as *O*^*t*^. The global patterning dynamics then follows *X*^*t*+1^ = *F*(*X*^*t*^, *O*^*t*^) with a global update function *F*.

We consider a patterning system of this type to be completely programmable, if (i) the organizer cells can steer any initial pattern *X*^0^ towards any desired target pattern *Y* in a finite time with a suitable time-dependent organizer input *O*^*t*^, and (ii) the time needed to reach the target pattern scales at most polynomially with the system size *L*. For instance, if the time to reach the target pattern increases linearly with *L*, we say that the system is completely programmable in linear time. In contrast, if the system would randomly generate different patterns, the expected time to produce a specified target pattern would increase exponentially with *L*. If complete programmability is not obtainable, we also consider partial programmability, where only a subset of target patterns is reachable. In either case, the sequence of organizer inputs, *O*^*t*^, may depend on the initial pattern *X*^0^. We first focus on the question of whether target patterns can be reached and will then consider the issue of stabilizing target patterns.

Note that while some cellular automata, including the paradigmatic ‘Game of Life’ introduced by Conway^17^, are well known to be universal computing devices, the question of programmable pattern formation is distinct from universal computing: In the context of computing, both the ‘program’ and the ‘input data’ are specified by the initial state of the CA. In contrast, for programmable pattern formation the initial state is arbitrary, while the target pattern is encoded in the state transitions of the organizer cells, and the patterning algorithm is specified by the update rule and the topology of the system.

### Some update rules enable complete programmability

The discrete model facilitates an efficient computational test for complete programmability, which relies on the representation of the patterning dynamics by a directed graph (Fig. 2a-c). In this ‘patterning graph’, each patterning state of the system is represented by a node. A connecting arrow from node *X* to node *Y* indicates that an input signal *B* exists that takes the system from state *X* to state *Y* in one step, i.e., *Y* = *F*(*X*, *B*). The arrow is labeled with this input signal *B* (if multiple signals exist, the arrow gets multiple labels). Complete programmability is then equivalent to the statement that there is a path from every node to every other node in the patterning graph, i.e., that the graph is strongly connected^18^ (Fig. 2). Strong connectivity of directed graphs with tens of thousands of nodes can be rapidly tested with standard algorithms (Suppl. Notes S1.1).

**Figure 2.**
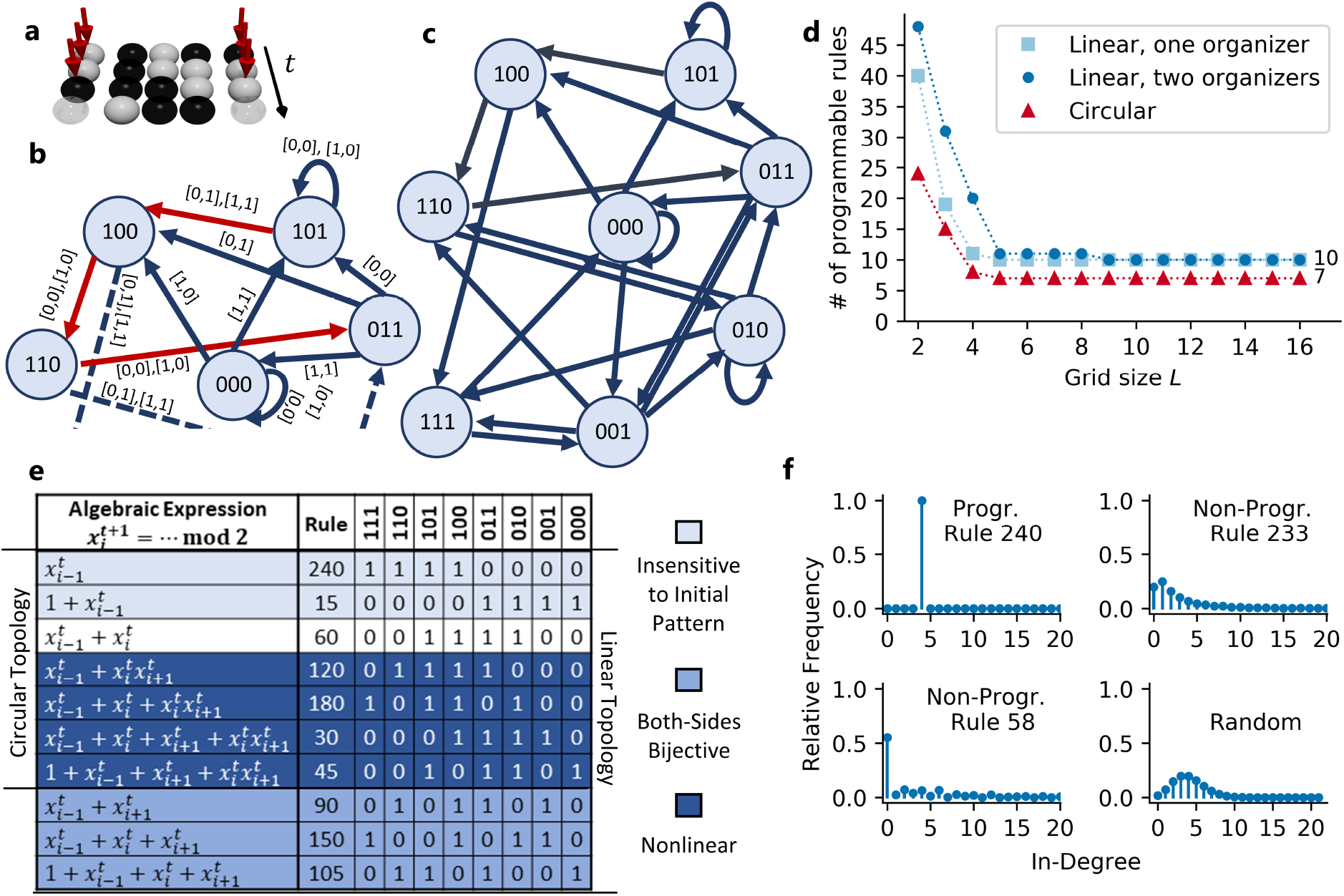
Patterning graph. **(a)** Illustration of the patterning dynamics of a small system (*L* = 3), steered from initial pattern 101 to target pattern 011 with two-sided control under rule 86 (cf. Fig. 1). **(b)** Part of the corresponding patterning graph, with nodes representing patterns and arrows possible transitions. The organizer inputs that can trigger a transition are indicated next to the corresponding arrow, e.g., 0,0 when the left and right organizer cell both supply a ‘0’ signal (sometimes multiple input combinations are possible). The path corresponding to (a) is highlighted in red. **(c)** A system with a given update rule permits programmable pattern formation, if the full patterning graph is strongly connected, i.e., there is a path from every node to every other node (regardless of the required inputs). **(d)** The number of distinct programmable rules decreases monotonically with the system size and reaches a plateau-value (10 for linear topology, 7 for circular topology). **(e)** The update rules that remain programmable for all system sizes with circular (first 7) and linear topology (all shown rules), listed in their algebraic form, decimal rule number, and update map. Different shades indicate properties of the rules that are discussed in the main text. **(f)** Statistical characterization of patterning graphs by their in-degree distribution (linear systems with two organizer cells, *L* = 16 bulk cells, and three different exemplary rules; distributions shown up to in-degree 20). All programmable rules have the same distribution as the shown rule 240, whereas the distribution of non-programmable rules is broad (shown examples: rule 233 and 58), except for the *identity* and *complement* rule (distributions for all rules shown in Fig. S2). For comparison, the in-degree distribution of a randomized graph is also shown (random redirection of arrows to any node with equal probability).

We applied this test in systems with different sizes and topologies, for the minimal model of *k* = *2* states, where the number of possible update functions *f* is only 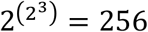 Given that our underlying model is symmetric with respect to the spatial directions ‘left’ and ‘right’, and also with respect to the internal states ‘0’ and ‘1’, the set of 256 rules can be split into 88 equivalence classes, which we refer to as ‘distinct rules’. We tested all rules and found that there are several which enable complete programmability of pattern formation. The number of such rules depends on the topology and decreases with the system size *L* (Fig. 2d). Strikingly, however, for systems of size *L* ≥ 9 the numbers no longer decrease, and the set of remaining rules is unchanged up to the maximum size that we were able to test numerically. This observation suggests that a subset of ‘programmable rules’ enables complete programmability of pattern formation for systems with a fixed topology but any size. This subset contains ten distinct rules for linear topology and seven distinct rules for circular topology (Fig. 2d,e). In the linear topology, the optimal placement of organizer cell(s) with respect to the number of programmable rules is at the boundaries (Suppl. Notes S1.2 and Fig. S1). The ten distinct rules for linear topology were also independently^19^ identified by another study^20,21^.

While complete programmability, as defined above, is a global property of the patterning graph (strong connectivity), the ten rules of Fig. 2e also stand out in a local property of the patterning graph, the in-degree distribution. In a directed graph, the in-degree of a node corresponds to the number of incoming arrows. Fig. 2f shows histograms of the in-degrees of all nodes in the patterning graphs of one programmable and two non-programmable rules. For comparison, Fig. 2f also shows the in-degree distribution of a randomized graph, in which the outgoing links from each node are randomly reassigned to any target node. These examples illustrate an empirical property (Fig. S2 and Suppl. Notes S2): Whereas programmable rules have an in-degree distribution with only a single peak at in-degree two (for one organizer cell) or four (for two organizer cells), non-programmable rules display a broad in-degree distribution, which depends on the specific rule and differs from the distribution for a randomized graph. This indicates that one can distinguish programmable from non-programmable rules already based on the local structural statistics of the patterning graph, which for large systems could also be sampled from a randomly chosen subset of nodes. We will see further below that the number of programmable rules increases rapidly with the number *k* of states.

### Different mechanisms for complete programmability of patterning

To illustrate the mechanisms by which complete programmability is achieved, Fig. 3 displays the spatio-temporal dynamics for several rules, topologies, and target patterns. The simplest mechanism is that of Fig. 3a, where a single organizer cell feeds its changing state into a linear array of bulk cells, in this case from the left edge. The bulk cells propagate the received information to their neighbors, effectively writing a temporal signal into a spatial pattern. This mechanism is reminiscent of the clock-and-wavefront mechanism in vertebrate somitogenesis, which relies on a temporal oscillation that is converted into a spatial stripe pattern^22–25^.

**Figure 3.**
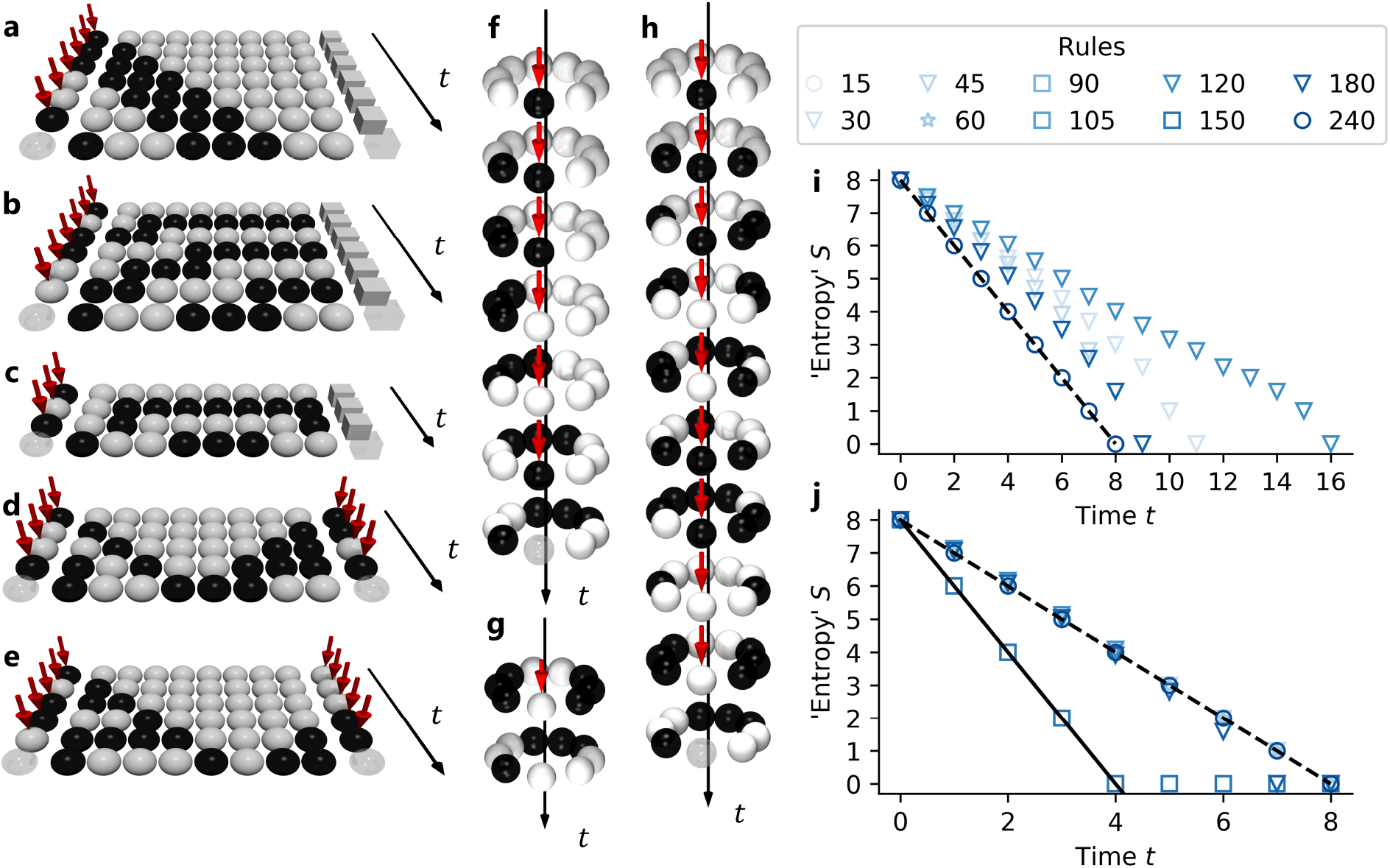
Patterning dynamics. Exemplary kymographs show the patterning dynamics of different systems, each with *L* = 8 bulk cells, but different cell arrangements and update rules: **(a)** linear system with one organizer cell and rule 240, **(b)** rule 15, and **(c)** rule 105, **(d)** linear system with two organizer cells with rule 90, **(e)** two-sided control with rule 30 and a different target pattern, **(f)** circular system with embedded organizer cell and rule 240, **(g)** rule 30 starting from a non-homogeneous initial state, and **(h)** from a homogeneous initial state. **(i)** Characterization of the patterning dynamics by the entropy-like observable *S*(*t*), a logarithmic measure of the number of different patterns that remain after(*t*) update steps, if the patterning process is started from the ensemble of all possible initial states (see main text). The data shows the time-dependence of *S*(*t*) for different update rules (symbols) in a circular topology with the same target pattern as in (f). The dashed line marks *S* = −*t* + *L* for comparison. The symbols for the rules are chosen according to the four categories introduced in Fig. 2e. **(j)** As in (i), but for linear topology with two organizers. The solid line additionally marks *S* = −2*t* + *L* for comparison. The behavior shown for this particular target pattern is generic, as can be seen from Figs. S3 – S5, which show the minimum, maximum and average *S(t)* over all patterns.

The examples in Fig. 3b-e display more complex behavior, suggesting alternative modes of programmable pattern formation. Fig. 3b displays a variant of the mechanism in Fig. 3a, which updates a cell to the inverted state of its left neighbor. Thereby, this rule produces a dynamics where the pattern oscillates as it is pushed from the organizer cell into the bulk. In both cases, the initial state of the system is completely erased during the patterning process, such that the sequence *O*^*t*^ of organizer inputs is independent of the initial pattern *X*^0^. In contrast, a third rule (Fig. 3c) generates the same target pattern partially from the initial state, exploiting the computational power of the update rule. As a consequence, the target pattern is reached more rapidly. Figs. 3d and e illustrate programmable pattern formation with simultaneous input from two organizer cells. Only update rules that are affected by input signals from both the left and the right side can simultaneously process information from two organizer cells. In the case of Fig. 3d, rule 90 produces the target pattern by symmetrically using information from both organizer cells, while Fig. 3e illustrates an asymmetric pattern formation process with rule 30, where information from the left side is preferentially used. Finally, Fig. 3f-h display kymographs for cases where a single organizer cell is embedded in a ring of bulk cells. In Fig. 3f, the information from the organizer cell is pushed only in one direction, such that the patterning process is analogous to that of Fig. 3a for the linear cell array. In contrast, Fig. 3g illustrates a case where the target pattern is computed (by rule 30) from the initial pattern. The most complex example is that of Fig. 3h, where rule 30 propagates information from the organizer cell to both sides, producing an ‘interference’ phenomenon when the two signals meet, which results in a much longer time required to reach the target pattern.

The unifying principle underlying these examples is linear transport of patterning information from one or multiple sources, with concurrent processing of this information by the bulk cells. The behavior is visually simple only if the bulk cells merely pass on the information they receive, while additional signal integration with their internal states typically generates complex spatio-temporal dynamics.

### Time to reach the target pattern

The examples in Fig. 3 suggest that systems with programmable rules are completely programmable in linear time. With simple unidirectional transport of patterning information (as in Fig. 3a, b, and f), the maximum number of update steps in an optimal path is equal to the number *L* of bulk cells. With two organizer cells, this maximum can be cut in half, as seen in Fig. 3d. Furthermore, in some cases the update rule can construct a portion of the target pattern from the initial pattern to speed up the pattern formation process (Fig. 3c and g).

We obtain a global view of the patterning dynamics by considering the ensemble of all possible initial states of the system and monitoring how this ensemble progressively shrinks towards a single point in state space (the target pattern). A convenient observable to characterize these dynamics is the time-dependent “entropy” *S(t)* = log_2_(Ω(t)), where Ω(t) denotes the number of points in state space occupied by the ensemble at time *t*. As the patterning process proceeds from every possible initial state along every possible shortest path to the target state, *S*(*t*) decreases from *L* to zero. The computed time traces *S*(*t*) for a system of size *L* = 8 with the different programmable rules are shown in Fig. 3i (circular topology) and Fig. 3j (linear topology with organizer cells at both ends). In the latter case, *S*(*t*) decreases roughly linearly for all rules, corresponding to an exponentially shrinking volume of the pattern ensemble in state space. The velocity of this “entropy reduction” is either 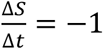 or −2 (dashed and solid line, respectively). In the circular topology, *S*(*t*) either decreases linearly with slope −1, or displays a slower decrease with variable slope. Taken together, the dynamics of *S*(*t*) is consistent with the spectrum of behaviors observed in the examples of Fig. 3a-h. It is also consistent with the behavior of the average shortest path length in the patterning graph (Suppl. Notes S3).

### Conservation principle and programmable rules for *k*-state systems

Intuitively it is clear that faithful transport of patterning information from organizer cells into the bulk requires a conservation principle. This notion is formalized by the concept of bijectivity. We define a rule *f* to be left-bijective, if the mapping *x* → *y* with 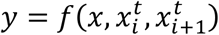 is bijective for each combination of 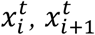 values (Fig. 4a,b). For a left-bijective rule every possible output 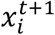 can be reached by choosing an appropriate left input 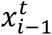, irrespective of 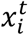 and 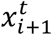 This property suffices to guarantee that one can find a series of inputs *O*^*t*^ from an organizer cell on the left to produce any target pattern in the bulk cell array (Fig. 4c and Suppl. Notes S1.3, S1.4). Similarly, if an update rule is right-bijective, it permits complete programmability of pattern formation from an organizer cell on the right. The argument of Fig. 4c is constructive in the sense that it not only guarantees the existence of a suitable organizer sequence *O*^*t*^ to reach the target pattern, but it provides a recipe to explicitly construct *O*^*t*^, given the update rule as well as the initial and the target pattern (Suppl. Notes S1.5). This recipe confirms the distinction between the simple rules of Fig. 3a, b and the other programmable rules with more complex behavior: For the simple rules, the organizer sequences *O*^*t*^ can be chosen independent of the initial state of the system, whereas for the complex rules, the construction of *O*^*t*^ requires knowledge of the initial state (see the classification of rules in Fig. 2e). Rules that are both left- and right-bijective can faithfully transport information from both sides, which can speed up the patterning process with two organizer cells, as seen in Fig. 3d and e. However, in the circular topology, rules that are both left- and right-bijective do not enable complete programmability, since they are unable to convert initially symmetric patterns into an asymmetric one (Suppl. Notes S1.6).

**Figure 4:**
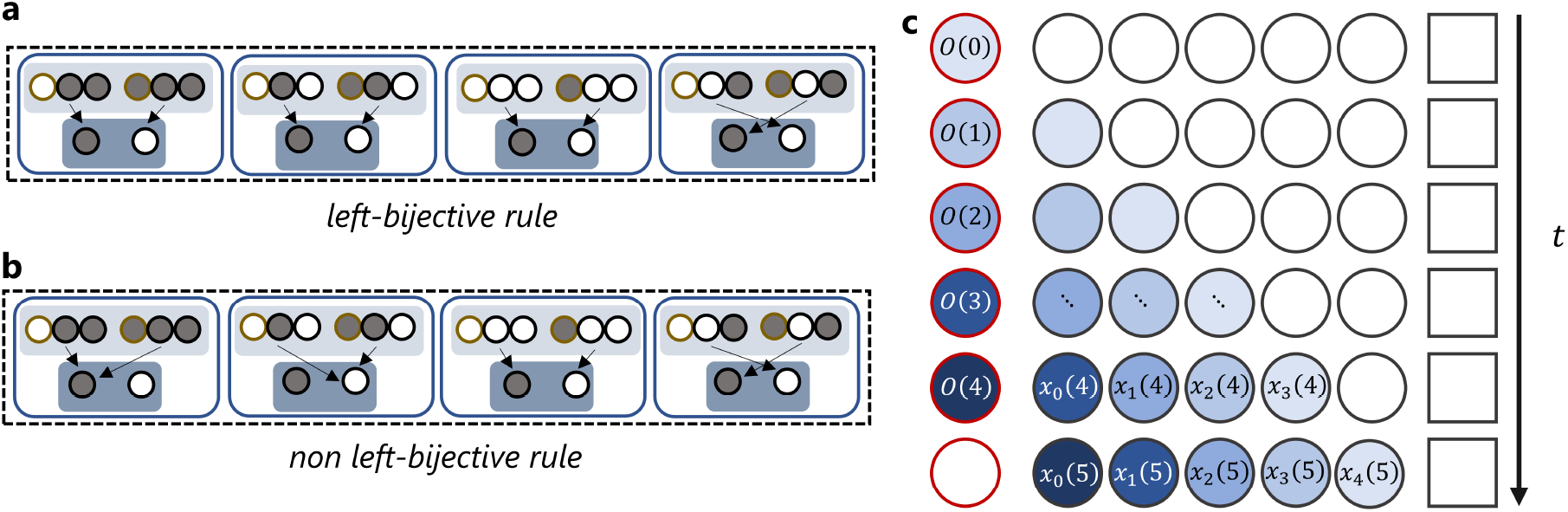
Construction of the organizer sequence for a left-bijective update rule. **(a)** Illustration of a left-bijective rule for an elementary two-state cellular automaton. The eight possible input configurations are grouped into four pairs (light blue background) according to the states of their middle and right-hand cells (dark boundaries). For each of these pairs, the rule establishes a one-to-one mapping between the state of its left input cell (light boundary) and its output (dark blue background). Fig. S6 illustrates the bijectivity of all programmable rules in a similar way. **(b)** In contrast, a rule that maps at least one pair of input states onto the same output state is not left-bijective (first and second pair in the shown example). **(c)** Illustration of the construction scheme for the organizer sequence *O*(*t*) that steers an initial pattern to a desired target pattern. In this example, the system has *L* = 5 bulk cells and an organizer cell on the left (red boundary). The construction scheme determines the organizer sequence by backward propagation from the target pattern, and explicitly demonstrates that bijectivity implies programmability. All white cells are not influenced by the organizer sequence *O*(*t*), so their states can be computed from the initial pattern with the update rule. Back propagation then begins by setting the value of the rightmost cell in the final pattern to the desired value *x*_*i*_(5). Since *x*_*i*_(4) and *x*_*i*_(4) are known, left-bijectivity guarantees that there is a value *x*_*i*_(4) such that *x*_*i*_(5) has the desired value. Similarly, it is possible to set *x*_*i*_(3) such that *x*_*i*_(4) takes on the required value determined in the previous step. Iterating along the diagonal with the lightest blue shade then fixes the first organizer input, *O*(0). Back propagation of *x*_*i*_(5) then fixes *O*(1) and so forth. Thus, bijectivity of the update rule suffices to construct an organizer sequence *O*(*t*) to steer a given initial pattern into any desired target pattern.

Importantly, the argument of Fig. 4c is valid for any length *L* of the system, for any number *k* of cell states, and it can also be generalized to obtain a constructive recipe for update rules that depend on larger neighborhoods of cells (Suppl. Notes S1.7). The bijectivity property can be used to show that with *k* cell states there are at least 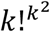 programmable rules before taking into account symmetries, but that the fraction of bijective rules within all rules decreases rapidly with *k* (Suppl. Notes S1.8). Furthermore, it follows (Suppl. Notes S1.9) that the maximal length of the shortest path from a given initial pattern to a desired target pattern is *L* in the case of one organizer cell, and 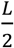 for two organizer cells and both left- and right-bijective rules (or 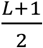 when 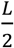 is not an integer), as we had empirically seen above.

### Robustness against errors and error correction

In real systems, the communication between subunits, as well as the information processing within subunits, are exposed to noise, causing some level of stochasticity in the dynamics. How sensitive programmable pattern formation is to such stochasticity is therefore a crucial question. To explore this question, we extend our model by introducing an error process. After executing the deterministic update rule, each cell stochastically switches to the opposite state with probability *p*, or remains in its state with probability 1 − *p*, independent of the state of its neighbors. This stochastic update models effects such as loss of memory (of the prior cell state), noise in the internal regulatory circuit that encodes the update rule, unreliable signal transmission from neighboring cells, and noise in the exact timing of state transitions. Since we consider a spatially and temporally homogenous system, we take *p* to be constant in time and space.

To monitor the impact of the stochasticity, we measure the reliability of the pattern formation process as a function of *p*. Specifically, we determine the probability that the final pattern has no error, i.e., that it matches the desired target pattern (Suppl. Notes S4.1). For programmable rules that take input from only one neighboring cell, this probability can be estimated as

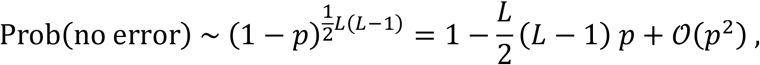

since a system of size *L* reaches its target state after at most *L* steps, and errors in *L*(*L* − 1)/2 individual cell updates can have an influence on the final pattern (see Fig. 4). This error estimate helps to interpret our simulations of the model (Fig. 5). The reliability as a function of the error rate *p* in a system of fixed size is shown in Fig. 5c for all programmable rules (red symbols), confirming that the estimate (solid red line) captures the essential behavior of the model. In particular, the reliability decreases linearly with *p* for small error rates (dashed red line), implying that all patterning schemes considered so far are very sensitive to errors.

**Figure 5:**
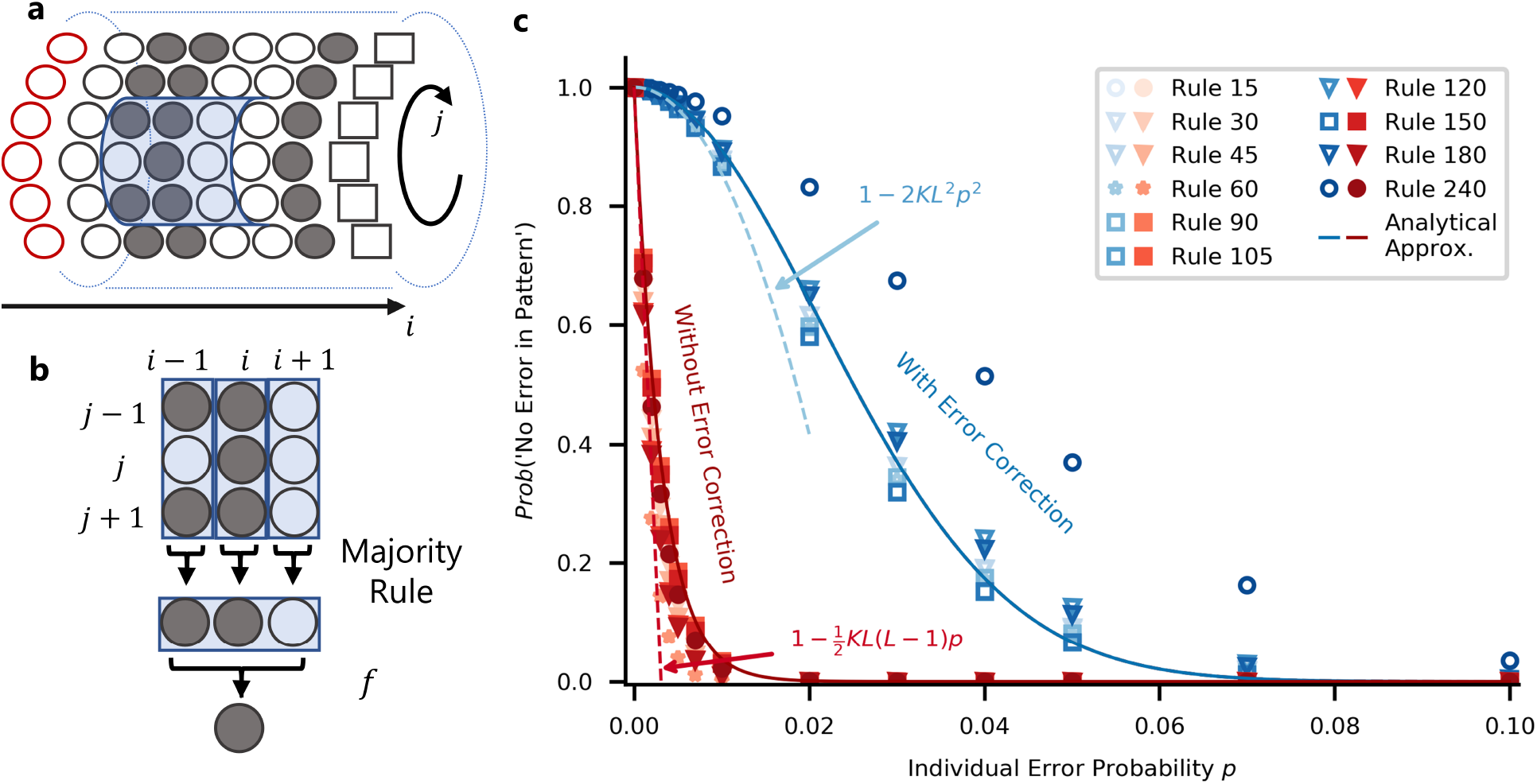
Robustness against errors and error correction. **(a)** To explore a mechanism for error correction, we consider a cylindrical system with an array of organizer cells along the left edge and a fixed boundary condition along the right edge. The update rule now takes input from a 9-cell neighborhood (shaded blue). **(b)** Given a 9-cell neighborhood, the update rule applies “majority voting” in the vertical direction (index *j*), establishing a consensus triplet to which one of the 1D programmable rules *f* is applied, yielding the final output. **(c)** Computer simulations (symbols) and analytical theory (lines) for systems with error correction by majority voting (red) compared to the case without error correction (blue). As in Fig. 3, the symbols for different rules are chosen according to the four categories introduced in Fig. 2e. In each case, a system of size *L* = *k* = 9 was used. Sufficient simulation runs were performed to estimate the plotted probability of arriving at the correct final pattern with a statistical error smaller than the symbol size (see Suppl. Text S4). Convergence is demonstrated by the observation that rules for which the same error behavior is expected (Rules 15 and 240) yield data points lying on top of each other. See Suppl. Text S4 for the analytical approximations.

The root cause of the high sensitivity to errors is the one-dimensional geometry of our model systems: A single failure breaks the “chain of command” from the organizer cells to the distant bulk cells. Given that most real systems have two- or three-dimensional arrangements of subunits, it is natural to extend the spatial dimension of our model. We focus on a two-dimensional extension of our model, in which *K* parallel cell lanes, each of length *L*, are connected to form a tube (see Fig. 5a, where periodic boundary conditions are applied in the vertical direction). The parallel lanes offer redundancy, which the cells can leverage to increase the reliability: They communicate with their lateral neighbors and apply a majority voting rule for their update (Fig. 5b), which in tissues could be mediated by diffusible signaling molecules. Along the axis of the tube cells follow the same rules as in the one-dimensional model above. The blue symbols in Fig. 5c show the reliability as a function of *p* for a tube with the same length *L* as the one-dimensional system (see caption for parameters and Suppl. Notes S4.1 for the numerical procedure). We observe a dramatic increase in reliability, caused by the ability of the lateral majority voting rule to correct isolated errors. The error correction changes the scaling of the reliability with *p* from linear to quadratic (dashed blue line). In fact, the observed behavior can be captured by an estimate (solid blue line) based on counting the number of arrangements of errors that cannot be corrected (Suppl. Notes S4.2). The expansion of this estimate for small *p* shows that

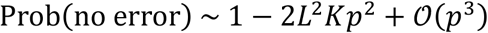

for the tube. The rules which only shift the state of the cell to the next cell perform best, since they spread errors the least (Suppl. Notes S4.3).

### Robustness against variable timing of organizer signals

The above analysis showed that local organizers can steer the bulk cells into any one-dimensional target pattern using only local signals processed according to simple rules. However, this requires precise timing in the switching of the organizer signals. Precise timing is also needed for the arrest of the patterning process when the target pattern is reached, because the target pattern is generally not a fixed point of the dynamics (programmable rules have only trivial stationary patterns). To explore the degree of programmability that can be achieved with less precise timing, we consider an alternative scheme, which uses update rules with nontrivial stationary patterns: For each organizer input, we let the system evolve until the pattern no longer changes before applying a new input. Together with each input, we also allow a global change of the update rule (same for all cells). In a developmental system, this would amount to a change in the interpretation of intercellular signals in different developmental stages, which is a known phenomenon, e.g. for the Toll signaling pathway of Drosophila^1^. The change could be triggered by a global signal, which does not need to be timed precisely, since the system runs into a stationary pattern at which it can stay for an extended time. Global changes of the update rule could in principle also be implemented in a synthetic DNA-based system (Suppl. Notes S5).

For simplicity, we refer to the combination of an input with an update rule as an ‘instruction’. We only use instructions that lead the system to a stationary state, avoiding those that lead to limit cycles. To construct an efficient search method for a protocol that steers the system from a given initial pattern to a desired target pattern, we first analyze the patterning graphs of all CA rules. For each rule and organizer input, we identify all attractors and their basins of attraction, which consist of all configurations from which the attractor is reachable (Fig. 6a). We then construct a single ‘attractor graph’ from all basins of attraction, by adding a directed link *X* → *Y* for each pattern *X* in the attraction basin of pattern *Y*. Each link has an associated instruction. Using the attractor graph, we determine the instruction sequence by extracting the shortest path connecting two patterns (Suppl. Notes S3). This recipe minimizes the number of instructions, but other objective functions, such as minimizing the number of changes in the rule or the total time needed to reach the final attractor, could be implemented with similar methods.

**Figure 6:**
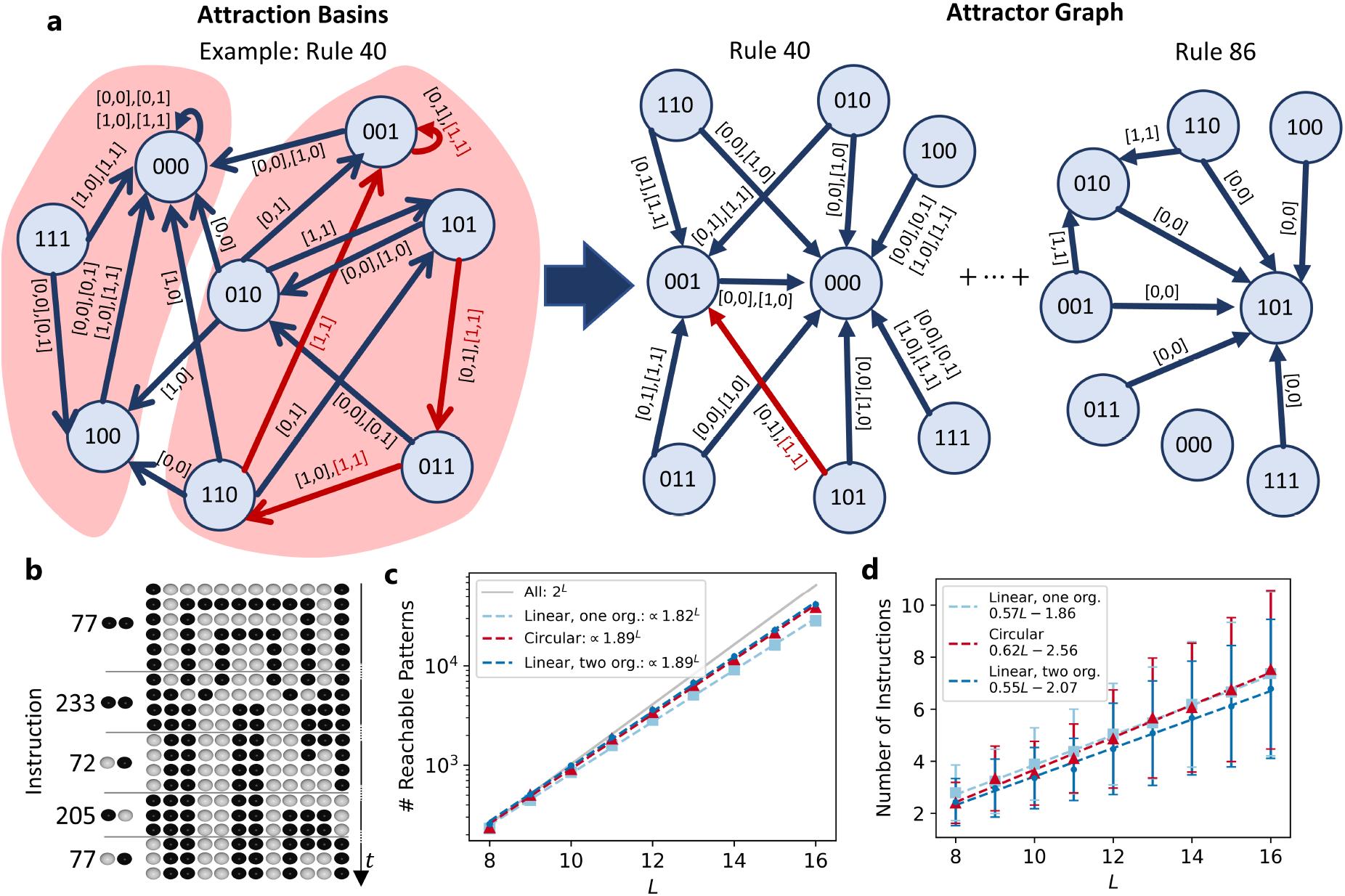
Robustness against variable timing of organizer signals. **(a)** To explore a mechanism for programmable pattern formation that is robust against variable timing of organizer signals, we consider an alternative scheme for programmable pattern formation, which uses all update rules with nontrivial stationary patterns (see main text). Left: Example of a patterning graph of rule 40 with *L* = 3 in a linear topology with two organizer cells. In dark red a sample path is shown leading from pattern ‘101’ to the fixed point ‘001’ using the instruction (rule 40, [1, 1]) – i.e., left and right organizer cell have both state 1. The light red shaded areas show the attraction basins of the instruction (rule 40, [1,1]) with the attractors ‘000’ and ‘001’. Using also the other possible instructions with rule 40 all configurations are in the attraction basin of ‘000’, while only the right shaded subset is in the attraction basin of ‘001’. Right: The contributions of rule 40 with all possible inputs to the attractor graph are calculated by adding a directed edge towards a node if the pattern corresponding to the origin of the arrow is in the attraction basin of the target node. The red arrow corresponds to the red path on the left. The other contributions shown are from rule 86 depicted in Fig. 2. All contributions from all rules generate the attractor graph. **(b)** Example for *L* = 10. The target pattern is reached with 5 instructions. After each instruction, the system is allowed to reach its steady state which may last as long as desired (dots in timeline). **(c**) Number of reachable patterns as a function of *L* compared to the total number of patterns *2*^*L*^. The number of reachable patterns is determined by calculating how many nodes (patterns) of the attractor graph can be reached on a directed path from the 0 pattern. **(d)** Average number of instructions necessary to generate reachable patterns, with standard deviation as error bars, as a function of *L*, with a linear fit in the range *L* ∈ 8, 16 to exclude finite size effects.

We only consider the homogeneous initial condition (all cells in state ‘0’) with no prior spatial information that could seed the generation of patterns. Not all patterns can be reached by this scheme for larger system sizes (Suppl. Notes S6.1). We performed an exhaustive analysis to determine the reachable patterns for grid sizes up to length *L* = 16. We fit the resulting data in the saturated range *L* ∈ 8, 16 and determined the number of reachable patterns to scale as 1.89^*L*^ (circular and linear topology with two organizer cells) and 1.82^*L*^ (linear topology with one organizer cell), showing that even if not all patterns are reachable, an exponentially growing number is (Fig. 6c). Interestingly, approximately the same scaling applies for the linear topology if we weaken the assumption that the cellular automata can distinguish left from right, i.e., include only outer-totalistic rules in the attractor graph, which are agnostic to the directionality of the signals (Suppl. Notes S6.2). These empirical observations are consistent with the results of an analytical approach to determine the number of attractors of finite CA, which also indicates that the number of attractors for individual rules (except the identity rule 204) grows more slowly with *L* than the total number *2*^*L*^ of possible patterns^26^.

To characterize the patterning dynamics, we calculated the average shortest path length in the attractor graph, i.e., the average minimal number of required instructions, to reach the accessible target patterns as a function of the system size *L*. The empirically observed linear dependence (Fig. 6d) indicates that, even as the number of reachable patterns increases exponentially, the time, measured in number of instructions, to reach a target pattern increases only linearly with system size, as in our original scheme for programmable patterning (Fig. 3i,j).

Models for pattern formation processes can also be regarded as a means to compress the information required to specify a pattern. This notion is formalized by the concept of Kolmogorov complexity of a pattern, defined as the length of the shortest program for a Turing machine which outputs that pattern and halts^27^. Within our scheme, we can say that the complexity of a pattern is measured by the number of instructions needed to generate it starting from the homogeneous initial condition. Empirically, the patterns which require the fewest instructions exhibit some periodicity, which makes them amenable to compression, while there is no obvious visible difference between the most complex reachable patterns and the unreachable patterns (Suppl. Notes S6.3, Fig. S15).

## DISCUSSION

Programmable pattern formation in cellular systems is a remarkable phenomenon in biology, and a long-term goal for the design of synthetic multicellular systems^28^. Here, our objective was not to study any specific system, but to identify general schemes whereby local signals from organizer cells can direct global pattern formation. We chose a cellular automata-based modeling framework, which is sufficiently general to encompass a broad class of model systems, yet simple enough for explorative studies. For elementary cellular automata, in which cells have only two states and two neighbors, we performed an unbiased exhaustive analysis of all dynamical rules. We were then able to generalize some of our results to more complex systems. In particular, our approach led to the following findings: (i) Complete programmability of pattern formation by isolated organizer cells is possible only with a small fraction of distinct rules (Fig. 2), which fulfill a conservation principle for the transmission of patterning information (Fig. 4). (ii) While the detailed patterning dynamics implemented by each programmable rule is different (Fig. 3), the unifying principle can be interpreted as a generalization of the ‘clock-and-wavefront’ scheme underlying vertebrate somitogenesis, where a temporal signal is converted into a spatial pattern. (iii) Global pattern formation controlled by isolated locally acting organizers is intrinsically susceptible to errors, but the accuracy of pattern formation can be substantially improved with a simple error-correcting scheme based on local majority voting (Fig. 5). (iv) Programmable pattern formation controlled by organizer cells is generally sensitive to the timing of organizer inputs, but robustness against variable timing is achievable with organizers that have the additional ability to change the update rule of bulk cells, i.e., the way in which bulk cells interpret the received signals (Fig. 6).

Our results contribute towards a conceptual framework for constructing molecular or cellular systems with the ability of programmable pattern formation. DNA-based systems form a promising platform for molecular realizations of programmable pattern formation due to their programmability and information processing ability^29,30^. An elementary CA with the programmable rule 90 has already been implemented with DNA tiles^31,32^ albeit not in a way that permits inputs from an organizer. Other DNA-based implementations, which are more complex but offer more flexibility, have also been proposed^10^. On the basis of these proposed designs, a biomolecular CA that allows for input signals controlling its patterning process could be implemented as described in Suppl. Notes S5.

Synthetic cell-like systems with the capability to communicate and process information have also been implemented, based on emulsion droplets^9^ and liposomes^33^. Information processing within such synthetic cells is realized with artificial gene circuits, based on in vitro transcription or transcription-translation systems, whereas communication between neighboring cells is enabled, for instance, by dedicated protein pores^28^. A complementary path to achieve programmable pattern formation in cellular systems is to equip biological cells with engineered sensing and response systems^34,35^.

Given that our model was not designed to mimick any specific system, it is noteworthy that it led us to a principle of programmable pattern formation, which can be regarded as a generalization of the clock-and-wavefront scheme underlying vertebrate somitogenesis^22–24^. The basic principle is the same as that of a tape recorder, where a temporal audio signal is written into a spatial magnetic pattern. In the case of the clock-and-wavefront scheme, a periodic gene expression signal generates a stripe pattern via a determination front, which sweeps the tissue and arrests cells in their current state. The patterning dynamics displayed in Fig. 3 generalizes the clock-and- wavefront scheme by allowing for (i) any target pattern, not just regular stripes, and (ii) simultaneous transport and processing of patterning information. While our model does not implement a determination front, we considered an alternative scheme, in which the update rule of the bulk cells is also controlled by the organizers and the target pattern is stabilized dynamically. CA with changing update rules are interesting also from the computational perspective, since they were previously found to display capabilities linked to the computational problem of open-ended evolution^36^.

The simplicity of our model was key to obtaining rigorous results, but also poses limitations. One important limitation is the restriction to discrete internal cell states. Pattern formation processes are typically described by nonlinear dynamical systems with multistable behavior, such that qualitatively distinct patterning states, e.g., gene expression ‘on’ or ‘off’, can emerge in spatially adjacent regions. Our model adopts a coarse-grained level of description, which already assumes the existence of such discrete states and ignores all intermediate states. For a biological system, discrete update rules represent logic-based models of a biochemical signaling network^37^. For other types of systems, discrete update rules typically also represent ‘digital’ approximations of the underlying ‘analog’ dynamics.

Another limitation is the one-dimensional arrangement of cells within our model, which permits only linear propagation of patterning information in space. This restriction is somewhat relaxed in our quasi-1D extension of the model (Fig. 5), where lateral signaling between cells is used for error correction. However, this extension does not address the more general question of programmable pattern formation in two or more dimensions, which remains open for future work.

We also did not include biological processes like cell growth, cell division and death, but assumed that the patterning process occurs in a group of cells with a static arrangement, as for example in *caenorhabditis* vulva development^15^. Finally, we limited our study to a patterning scenario based on organizer cells. However, we found that the dynamical update rules of the bulk cells can also generate parts of the target pattern (Fig. 3), and considered an alternative scheme for programmable pattern formation, which combines patterning information from local organizers with “distributed computation” of patterning information (Fig. 6). Bulk cells with more states, or larger neighborhoods for the update rule, will have more computational ability and will therefore enhance the potential for programmable pattern formation via distributed computation. Indeed, it is well known that cellular automata can serve as computing devices, with some even shown to be computationally universal^38^. In those cases, the initial state of the cellular automaton serves both as the program and the input data, while the update rule specifies the mechanism of the computer and the result of the computation is obtained from the state after time evolution. The situation is different for programmable pattern formation: In our “organizer scenario”, the initial state of the system can be simple, e.g., homogeneous, while the input data (patterning information) is supplied as a time-dependent local signal. Bijective update rules enable universal pattern formation with this scenario. Interestingly, these bijective rules were among the “illegal” rules excluded in Wolfram’s pioneering study on the statistical mechanics of cellular automata^11^, due to their violation of the quiescence and isotropy conditions.

Our question of programmability is closely related to the question of ‘controllability’ in the field of control theory. Control theory provides a general mathematical framework to analyze the control of dynamical systems^39^. It formalizes the intuitive notion of ‘controllability’ as the ability to steer a dynamical system to any desired state from any initial state by appropriate signals. A focus of recent research has been on the control of complex networks^40,41^, a broad class of dynamical systems ranging from networks of protein interactions^42^ and neurons^43^ to power grids^44^. Application of control theory concepts to linear dynamics on networks with complex topologies led to insights about the relation between network topology and the controllability of its dynamics^40,41^. Here, we focused instead on systems with simple topologies but more complex dynamics and studied how the ability to control pattern formation depends on the dynamical rules that propagate patterning information into the system.

In conclusion, programmable pattern formation connects the experimental fields of synthetic and systems biology to theoretical research on self-organization, computation, and control. We established simple scenarios for programmable pattern formation in cellular systems based on local organizers. Our results provide a rigorous basis for the analysis of more complex patterning scenarios, and for a conceptual framework to design synthetic molecular and cellular systems.

## ACKNOWLEDGEMENTS

We are grateful to Kilian Vogele and Friedrich Simmel for helpful discussions. This work is supported by the German Research Foundation via the collaborative research center SFB1032 and the Excellence Cluster “ORIGINS” through UG.

## AUTHOR CONTRIBUTIONS

All authors designed the research and analyzed the data. T.R., S.K., H.W. performed simulations, T.R., S.K., U.G. wrote the paper.

## DATA AND CODE AVAILABILITY

The data that support the findings of this study, as well as the computer code used to generate this data, are available from the corresponding author upon reasonable request.

