## Supplementary Text and Figures for "Programmable pattern formation in cellular systems with local signaling"

### Supplementary Information

Tiago Ramalho[#], Stephan Kremser[#], Hao Wu, Ulrich Gerland*

Physics of Complex Biosystems, Physics Department, Technical University of Munich, James-Franck-Str. 1, D-85748 Garching, Germany

[#] These authors contributed equally.

### SUPPLEMENTARY NOTES

#### S1 Programmability

In this section we first describe how we computationally test for the complete programmability of an update rule (S1.1) and how the number of programmable rules depends on the organizer position (S1.2). Afterwards, we provide a more formal proof that bijective rules are programmable (S1.3 and S1.3). Then we analyze in more detail the complexity of the algorithm to construct the organizer sequence (S1.5) and discusses the programmability for the circular topology (S1.6). We generalize the argument for programmability for rules with larger neighborhood sizes (S1.7), describe how a lower bound for the number of programmable rules can be found (S1.8) and, finally, elaborate how long it takes at maximum to reach the target pattern using the construction algorithm for the organizer sequence discussed in the main text (S1.9).

#### S1.1 Computational Test for Programmability of an Update Rule

In order to test a rule for programmability we first have to construct the patterning graph. For a given rule and boundary states $B$, we perform one dynamical step for a pattern $X$, which takes the system to state $Y$. This corresponds to an arrow in the input graph from node $X$ to node $Y$. We now construct the input graph by creating a graph in the "python-igraph" library[1] and creating directed links between all nodes which are connected by one step in the dynamics. Having constructed the patterning graph, the igraph library provides a direct test if the graph is strongly connected. For exemplary cases, we also checked the results by performing a similar procedure with the computationally less efficient networkx library[2]

### S1.2 Optimal Positioning of Organizer Cells

A natural subsequent question is, if programmable pattern formation is also possible if the organizer cell sits between two sets of bulk cells which have fixed boundaries on the sides. We again performed an exhaustive analysis for total bulk sizes ranging up to 15 bulk cells. The number of programmable rules in dependence of the position of the organizer cell is depicted in Fig. Figure **S1**. Two main results can be drawn from these simulations. First, the number of programmable rules is maximal for the organizer cell being located at a boundary, i.e., the case analyzed above. Second, no rule permits programmability of the system at any position. Hence, we conclude that for practical cases the desired position of the organizer cell is at the boundaries and subsequently focus on these systems.

### S1.3 Proof of Programmability for Transitive Rules

Consider a CA system running on a finite lattice with length $L$. The space of all possible configurations -- its configuration space -- shall be denoted ${}^{L}\Omega$. Similarly, we will consider a CA system on an infinite lattice, with corresponding configuration space ${}^{\infty}\Omega$.

A CA rule is transitive if it can create a path from almost anywhere to almost anywhere in the space ${}^{L}\Omega$ [3]. Concretely, a rule is transitive if for any $X, Y$ in ${}^{\infty}\Omega$, any $\varepsilon, \delta > 0$, there exist $\tilde{X}, \tilde{Y}$ and $\tau$ with properties $\left|X - \tilde{X}\right| < \varepsilon$, $\left|Y - \tilde{Y}\right|$, $\tau$ in $\mathbb{N}$ such that $F^{\tau}\left(\tilde{X}\right) = \tilde{Y}$. Here, the norm is given by Tychonoff metric[4]

$$|X - Y| = \sum_{i=-\infty}^{\infty} \frac{|x_i - y_i|}{S^i}$$

Now suppose we have two finite configurations ${}^{L}X$and ${}^{L}Y$. Without loss of generality, we consider $L$ odd. Certainly, there are two arbitrary infinite configurations in ${}^{\infty}\Omega$ which contain the finite configurations in their center positions (i.e., the positions $[-l, \dots, 0, \dots, l]$ with $L = 2l + 1$. These restrictions shall be denoted ${}^{\infty}X|_L = {}^{L}X$, ${}^{\infty}Y|_L = {}^{L}Y$.

Now, let us set the constants $\varepsilon = \delta = \frac{1}{S^l}$. Using the property of transitivity, we know that there exist $\tilde{X}, \tilde{Y}$ and $\tau$ with properties $|X - \tilde{X}| < \frac{1}{S^l}$, $|Y - \tilde{Y}| < \frac{1}{S^l}$ such that $F^{\tau}(\tilde{X}) = \tilde{Y}$. Comparing with $|X - Y| = \sum_{i=-\infty}^{\infty} \frac{|x_i - y_i|}{S^i}$, this implies that the values of $X$ and $\tilde{X}$ in lattice positions with indices $[-l, \dots, 0, \dots l]$ must match exactly (and the same for $Y$ and $\tilde{Y}$). Formally, ${}^{\infty}X|_L = \widetilde{{}^{\infty}X}\big|_L$, ${}^{\infty}Y|_L = \widetilde{{}^{\infty}Y}\big|_L$

From the above discussion it is clear that there exist infinite configurations which obey $\widetilde{{}^{\infty}X}\big|_L = {}^{L}X$; $\widetilde{{}^{\infty}Y}\big|_L = {}^{L}Y$ such that $F^{\tau}(\tilde{X}) = \tilde{Y}$. Thus, there is a flow in configuration space which leads from an infinite configuration which contains ${}^{L}X$ to one which contains ${}^{L}Y$.

Having established that such a flow exists, how to induce it in a finite lattice? The only states in the infinite lattice which influence the dynamics in the finite central section are the leftmost and rightmost adjacent cells. Fixing the temporal evolution of those states in $F^{t}(\tilde{X})$ will completely determine the dynamics of the evolution of the finite, bounded system. We have thus found the desired organizer input and proven the programmability of any transitive rule for linear topology with two organizer cells.

If there is just one organizer cell in a linear topology (assume on the left side), the finite configuration is set up between $[-L, \dots, -1]$ with the cell at positition 0, the only thing that need to be modified to apply above proof is the definition of the Tychonoff metric to be

$$|X - Y| = \sum_{i=-\infty}^{0} \frac{|x_i - y_i|}{S^i}.$$

**S1.4 Permutive Rules**

It is not possible to determine *a priori* whether a rule is transitive or not. We can, however, show that the 10 rules considered above possess another property: permutivity[4].

A rule $f$ is called permutive if for fixed states $y$ and z, either the map $f(\cdot, y, z)$ (left-permutive) or $f(z, y, \cdot)$ (right-permutive) is bijective. The term permutivity or bijectivity can in our context be used interchangeably. While we find bijectivity to be more fitting in general, here we keep permutivity to adapt to the literature we are referencing. Out of the 256 elementary CA rules, 28 are permutive[5]: 15, 30, 45, 60, 75, 85, 86, 89, 90, 101, 102, 105, 106, 120, 135, 149, 150, 153, 154, 165, 166, 169, 170, 180, 195, 210, 225, 240. However, only 10 globally unique rules remain after taking symmetries into account: 15, 30, 45, 60, 90, 105, 120, 150, 180, 240. The concept of permutivity is most easily grasped by visualizing the transition tables which describe the rules[5]. Below, we will reproduce these transition tables for the 10 unique permutive rules and highlight permutivity of the rules (Figure S6).

It has been shown that permutive rules are transitive[4], and therefore programmable. Given that the rules found during the numerical exploration are permutive, the hypothesis that they are programmable for all $L$ is confirmed.

The bi-permutivity of rules 90, 105 and 150 implies that these rules can transmit information from both sides and is consistent with our observation that these are indeed the rules which can create patterns the fastest, with a maximum runtime of $\frac{L+1}{2}$ steps (see S1.9).

We can also use the permutivity to directly construct programmable rules for any number states $k$. In order to do so, the transition table has to be ordered analogously to Figure S6. Then, by connecting each possible input in a sub-box with one possible output in this sub-box such that there is a bijective mapping a permutive/bijective rule can be created.

### S1.5 Construction Algorithm for the Organizer Sequence $\boldsymbol{O}(\boldsymbol{t})$

From the argument that bijectivity of a rule is sufficient for complete programmability (compare main text Fig. 4), a straightforward algorithm for the construction of the organizer sequence $O(t)$ emerges:

1. Calculate the dynamics of the system above the 'diagonal' with values $x_i(i)$ (this dynamics is independent of the organizer sequence).
2. To set state $x_i(L)$ to $y_i$ try every possible of the $k$ states for $O(L-i-1)$ and calculate the dynamics of the cells $x_{i-j}(L-j)$ for $j \in \{0, \dots, j-1\}$ until the state $O_{L-i}$ that sets $x_i(L) = y_i$ is found. Do this for all $i \in \{L-1, L-2, \dots, 1{,}0\}$ (in this order).

The number of evaluations $n_1$ of the rule for step 1 is

$$n_1 = \sum_{i=1}^{L-1} (L-i) \overset{Gauss}{=} L(L-1) - \frac{L(L-1)}{2} = \frac{L(L-1)}{2}.$$

The maximum number of evaluations $n_{2,max}$ for step 2 is

$$n_{2,max} = k \sum_{i=1}^{L} i \overset{Gauss}{=} k\frac{L(L+1)}{2}$$

In total, there are at most

$$n_{max} = n_1 + n_{2,max} = k\frac{L(L+1)}{2} + \frac{L(L-1)}{2}$$

rule evaluations necessary to construct the input pattern, i.e., it grows linearly with the number of states $k$ and quadratically with the system length $L$.

The organizer sequence constructed by this algorithm will steer the dynamics to the desired target state in $L$ steps ($\left\lceil \frac{L}{2} \right\rceil$ steps for two-sided programmability and rules $90, 105, 150$ if the algorithm is applied from both sides and only half of the pattern is considered each time). Comparison with the results for the average path length, which is obtained when a patterning graph is used to construct the path and thus the organizer sequence, shows that in most cases the performance difference is mostly just a small constant (e.g., 1.6 steps on average for linear topology and one organizer cell, see S3) when finite size effects are neglected, i.e., a relatively vanishing contribution for larger grid sizes. Taking into account that the construction of a patterning graph is computationally expensive, scaling with $k^L$, compared to the $kL^2$ complexity here, a direct construction of the organizer sequence is computationally more efficient.

**S1.6 Programmability in Systems with Circular Topology**

For circular topology, the above proof cannot be applied directly since the organizer signals sent around in one direction interfere with the ones coming from the other direction. Thus, the proof can only be applied if the rule does not take into account the right or left neighbor, therefore from the prototypical rules only $15 \equiv (1+p)\text{mod}_2$, $60 \equiv (p+q)\text{mod}_2$ and $240 \equiv (p)\text{mod}_2$ are definitely programmable for arbitrary $L$.

There is also a simple argument why the otherwise always programmable, both sides bijective rules $90, 105, 150$ are not programmable for circular topology: The topology is completely left-right symmetric, regardless of the organizer input, and these rules are left-right symmetric. Thus, when having a left-right symmetric initial pattern it is impossible to break this symmetry and reach a pattern that does not have this symmetry. Hence, these rules cannot be programmable.

Now we want to give a more formal argument to support the above statement. Outer totalistic rules are intrinsically left-right symmetric, as well as the organizer cell positioning for the circular topology. Suppose that the initial pattern for the dynamics is also left-right symmetric (cp. Fig. SFigure **S7**), as well as the inputs (circular). Choose a cell in the grid at position $i$. Then the corresponding mirrored cell at position $L - i$ will have the same state. Their corresponding neighbors will also have the same state as the neighbors of cell $k$ but exchanged position, i.e., the cell at $i - 1$ $(i + 1)$ will have the same state as the cell at $L - i + 1$ $(L - i - 1)$. But as the rule is left-right symmetric, this will yield the same state for $i$ and $L - i$ in the next time step. Thus, the symmetry of the pattern is conserved. Programmability requires that from any initial pattern any other pattern is reachable, so for a programmable rule it should be possible to reach an asymmetric pattern starting from a symmetric pattern (regarding left-right symmetry). As this is not possible for outer totalistic rules they cannot be programmable.

Since we cannot apply the proof, we found for linear topology we can only rely on analyzing the patterning graph for the rule in question if we are interested in determining which rules are programmable for circular topology. An exhaustive analysis of the whole rule space is unfeasible for a number of states $k$ larger than 2. But from our exhaustive analysis of all rules with $k = 2$ we can guess that the most promising candidates for being programmable are rules which are bijective in one argument, but not the other. Even for $k = 3$ the number of rules with this property is too large to be tested exhaustively, but a sample random sample of 2000 rules being bijective in either their right or left argument was analyzed (cp. Fig. SFigure **S8**) yielding a fraction of programmable rules out of this subset of 17.8% for $L = 12$

**S1.7 Programmability for Cellular Automata with Larger Neighborhoods**

The proof can also be extended for cellular automata that do not depend on the nearest neighbors but potentially on some other neighbors. Then the necessary condition is that the rule is bijective in the leftmost (rightmost) argument to be programmable from the left (right) side, and that the number of organizer cells increase to the radius $r$ on that side of the rule, i.e. the distance the furthest cell on the left (right) side has to the cell that gets updated (cp. the example in Fig S **Figure**

**S9**). In principle, the same algorithm as before can be applied with the difference that in the first step not only the first cell but the first $r$ cells can be addressed and set to any desired value, in the second step then the next $r$ cells can be addressed, so that finally only $\left\lceil \frac{L}{r} \right\rceil$ steps are needed. Further, as shown in Fig. SFigure **S9**, caution must be taken to set the states of the organizer cells in the correct order. Since for using bijectivity the states of all other cells in the neighborhood must be known, the innermost organizer cell must be set first, then the next cell and so on. This principle must be obeyed at all times to ensure that bijectivity can always be used.

#### S1.8 Number of Programmable Rules for an Arbitrary Number of States

Since a rule that is bijective in one of their outer arguments must be programmable, a lower bound for the number of programmable rules $N_{\mathrm{PR}}$ can be calculated by calculating the number of rules that are bijective in their outer arguments. To this end consider w.l.o.g. a left bijective rule. This means, for fixed central and right cells, for any of the possible $k$ inputs into the left cell there must be a unique assignment of one of the $k$ outputs. There are $k!$ possibilities for doing this uniquely. The fixed central and right cell can each have $k$ possible values, thus in total $k^2$ combinations. For each of them the unique assignment of left input to output can be differently, accordingly there are in total

$$N_{\mathrm{LBR}} = (k!)^{k^2}$$

left bijective rules. For right bijective rules the argument holds of course as well, thus $N_{\mathrm{RBR}} = k!^{k^2}$. Some rules are bijective in both sides, therefore the number of bijective rules in total is bound by

$$(k!)^{k^2} \leq N_{\mathrm{BR}} \leq 2(k!)^{k^2}$$

For example, for the case of elementary cellular automata, $k = 2$, thus $(2!)^{2^2} \leq N_{\mathrm{BR}} \leq 2\ (2!)^{2^2}$, or $16 \leq N_{\mathrm{BR}} \leq 32$. The above numerically determined numbers were $N_{\mathrm{LBR}} = 16$ and $N_{\mathrm{BR}} = 28$.

Finally, this also gives us a lower bound for the number of programmable rules $N_{\mathrm{PR}}$,

$$(k!)^{k^2} \leq N_{\mathrm{PR}}.$$

To assess if it is numerically feasible to search for programmable rules by sampling the rule space and directly testing found rules for programmability (as described in S1.1) it is interesting to compare this lower bound with the total number of rules for a given number of states $k$,

$$N_{\mathrm{TR}} = (k)^{k^3}$$

Using Stirling's formula, the ratio of the total number of rules to this lower bound scales as

$$\frac{N_{\mathrm{PR}}}{N_{\mathrm{TR}}} = \frac{(k!)^{k^2}}{(k)^{k^3}} \sim e^{-k^3}$$

i.e., decreases exponentially. This indicates, that sampling the rule space becomes unfeasible as the number of states $k$ gets large and emphasize the usefulness of being able to construct programmable rules directly using the property that bijective rules are programmable (S1.4).

#### S1.9 Time to Reach Target Pattern

In the realm of finite discrete systems, it is not only interesting to show that any configuration can be reached, but also that the number of steps to reach the desired target pattern within the given dynamic is not in the order of the number of total patterns $O(k^L)$.

Following the argument for construction of the organizer sequence $O(t)$ in the main text and S1.5, for linear topology with one organizer cell the number of steps to reach the target pattern is at most $L \ll k^L$.

For organizer cells on both sides and both left- and right-bijective rules we can use inputs from the left and the right to inscribe the target pattern into the system. If the number of bulk cells is even, these signals sent from the left and the right do not interfere with each other until the target pattern is reached, since the pattern is constructed with the same speed as the signal is sent through the system which is the maximum speed of 1 cell per time step. Thus, it takes at most $\frac{L}{2}$ time steps to reach the final pattern. If the number of cells is odd, then it takes $\frac{(L-1)}{2}$ steps to inscribe the pattern into all cells but the one in the middle. To set this cell we consider one of the organizer cells as having a random but known state and set the state of other the organizer cell $O(0)$ accordingly to reach the target pattern (which is possible as guaranteed by bijectivity). Thus, we need $\frac{(L-1)}{2} + 1 = \frac{(L+1)}{2}$ steps in to total to reach the target pattern.

### S2 Local Funneling of the Patterning Dynamics

Intuitively it is clear that programmable rules should allow to reach many different states in just a few steps. If that is not possible it would be very likely that some states cannot be reached from a defined initial pattern. To quantify that idea, we look at the number of possibilities to reach a pattern. This is the number of arrows that point into a pattern (node), the 'in-degree'[6] (compare Figure S2 and Fig. 2f of the main text for the distributions of the in-degree). It includes the arrows coming from other nodes as well as arrows coming from itself and gives a measure how easy it is to reconstruct the dynamics, i.e., in how many possible patterns the system was a step before it reached the current state. If one normalizes the in-degree by the total number of arrows in the graph, then this represents probability $p_i$ to reach the target state $i$ from a randomly picked initial state in one step. The negative logarithm of this probability is then a measure how much the dynamics of the system is compressed when it moves into this state. Low values correspond to high compression, high values to low compression. To construct an observable for the whole rule the average of this quantity is taken, which resembles the well-known Shannon entropy:

$$H = -\sum_i p_i \log_2(p_i)$$

where the sum runs over all nodes of the graph. It can be interpreted as the number of bits that it takes on average to code a certain pattern in a set of patterns, where pattern $i$ appears with probability $p_i$ [7]. In our thought experiment the number of bits to characterize the starting node is equal to the length of the lattice since the node was chosen at random, so every node has equal probability of being chosen. But after one step the dynamics has driven the system with higher probability into nodes that have an in-degree above the average. That causes the entropy to decrease in general, since the probability distribution over the nodes becomes uneven. Thus, the dynamics reduces the unknown information in the system. We calculated the Shannon entropy for all rules and system topologies. For length $L = 16$ the results can be found in Fig. SFigure **S10**, which shows the entropy normalized to the length of the system $\frac{H}{L}$. It can be seen that only two kinds of rules have values $\frac{H}{L}$ close to 1: The programmable rules and the rules which can trivially be proven to be not programmable for the investigated topology. These are for all topologies $51$ and $204$, which are the $complement$ and the $identity$ rule and thus clearly not able to allow for a programmability of the patterning. Furthermore, for circular topology rule $90$, $150$ and their $0 \leftrightarrow 1$ complement are not programmable which can be excluded by symmetry arguments (see S1.6). Last for linear topology with one organizer on the left rule $102$ and $153$, which cannot be programmable because they do not depend on the side at which the organizer cell is located in this case. So, the mechanism by which the programmable rules allow the boundary cells to exert their control over the system is by keeping the dynamics of the system as unspecified as possible, i.e., the rule does not facilitate a certain prepatterning just due to its structure.

### S3 Shortest Path Length

In the main text and S1, we found that a certain set of rules permits programmable pattern formation. To structure the set of programmable rules we look at an observable that describes the pattern formation process for each rule on average for all initial and target patterns. The observable discussed in the main text was an entropic quantity. As a second observable the average number of steps to steer the dynamics from any initial to any final pattern if always the shortest possible path is taken is an interesting quantity. It is calculated with the igraph-library[1], which uses breadth first search for unweighted graphs as in our case, and gives a measure of the performance of the rule since it corresponds to the average shortest path length $\langle l_{\mathrm{path}} \rangle$ in the patterning graph. As a function of the number of bulk cells $L$, this can be approximated very well with a linear function $\langle l_{\mathrm{path}} \rangle = aL + b$ for $L \geq 8$, exemplary shown in Fig. SFigure **S11**. Then, depending on the topology, different classes appear:

- For an organizer cell on one side, all programmable rules have $\langle l_{\mathrm{path}} \rangle = L - 1.6$, i.e., they take on average 1.6 steps less than the maximal value $L$ to reach the target pattern. The offset 1.6 can be attributed to (partially) using the initial pattern to construct the target pattern.
- For organizer cells on both sides, three groups of rules appear: First, also rules with $\langle l_{\mathrm{path}} \rangle = L - 1.6$ appear, which have in common that they only take into account inputs from one side. Second, rules that take into account inputs from both sides in a bijective fashion have all $\langle l_{\mathrm{path}} \rangle = \frac{L}{2} - 0.34$. Third, rules that take into account inputs from one side in a bijective fashion and inputs from the other side in a non-bijective fashion lie in between the two former cases concerning the slope of the linear function.
- For circular topology, rules from the first and third group appear.

### S4 Robustness against errors

#### S4.1 Numerical Simulations

For a given rule and 1D initial and 1D target pattern of length $L$ the shortest path and its corresponding inputs are calculated. Then, for a given height $K$ of the 2D grid, the equivalent stripped system is simulated for a given $p$ with the 2D equivalent of the aforementioned rule, which is constructed as described in the main text. The final pattern is compared with the desired target pattern and, if they coincide, counted as having no error in the final pattern. This procedure is carried out $N_{\text{trial}}$ times for each possible combination of initial and target pattern.

In this way, the probability to get an error in the final pattern can be calculated as a function of the rule $R$, the grid dimensions $L$ and $K$, the single error probability $p$ and the number of trials $N_{\text{trial}}$ (if $N_{\text{trial}}$ is chosen large enough this dependency should be vanishing):

$$P_{\text{NoError}}(R, L, K, p, N_{\text{trial}}) \approx \frac{\#\text{ desired target} = \text{actual final state}}{\#\text{ trials to reach a target total}}$$

We found $N_{\text{trial}} = 5$ to be large enough to get convergence.

#### S4.2 Analytical Approximations of the Error Probability

To estimate the performance of this error correction mechanism it is worthwhile to investigate its properties more intensively, i.e., identify the error patterns that can be corrected and the patterns that cannot be corrected.

First the behavior in the bulk is considered, i.e., the effect of the boundary states is not considered, just the type of boundary present. Since the averaging process is independent in each vertical line, they can be considered independently. For such a line, possible successful and unsuccessful error corrections are shown in Fig. Figure **S12**. When the neighborhood slides over any three vertical cells it will produce an error in the majority determined if at least two of the three cells have the wrong state, see  Fig. Figure **S12**b,c, so in the next timepoint there is an error in this or one of the neighboring cells (in which exactly depends on the rule, see discussion below). But even if two errors are only one cell apart, this can be corrected for if no additional error is introduced in the next time step Fig. Figure **S12**b.

To derive the probability that such processes do not appear and lead to an error in the target pattern consider first a vertical lane with $K$ cells. Through the erroneous cellular automaton there shall be $0 \leq i \leq K$ cells with errors in the lane. First, compute the probability for no two errors with two or less non-erroneous cell between them. To that end the number of possibilities to arrange the errors in such a way is needed. This number can be calculated as follows. Each error gets assigned with two non-errors above it (cp. Fig. Figure **S13**). Then there are $i$ of these elements and $K - 3i$ non-errors left that can be ordered into the lane. There are

$$N_1 = \binom{K - 2i}{i}$$

possibilities of doing that. Of course, this excludes the case when the elements on the top of the lane is an error. For this case, the one error is fixed at the top and there are only $i - 1$ or $i - 2$ error elements and $K - i - (i - 1)$ or $K - i - 1 - (i - 1)$ non-error elements left to distribute, which can be done in

$$N_2 = \binom{K - 2i}{i - 1} + \binom{K - 2i + 1}{i - 1}$$

possibilities. In total, there are then $N_1 + N_2$ possibilities to have no two errors next to each other. Dividing this number by the number of total possibilities to distribute $i$ errors and $K - i$ non-errors,

$$N_{\text{tot}} = \binom{K}{i}$$

then gives the probability to have $d \geq 2$ correct cells between any two errors:

$$P_{1,d\geq 2}(m,i) = \frac{\binom{K-2i}{i} + \binom{K-2i}{i-1} + \binom{K-2i+1}{i-1}}{\binom{K}{i}}$$

The leading ‘1’ in the index indicates that only a single lane is considered.

For periodic boundary conditions the formula is accordingly

$$P_{1,d\geq 2,\mathrm{PB}}(m,i) = \frac{\binom{K-2i}{i} + 2\binom{K-2i-1}{i-1}}{\binom{K}{i}}$$

After having specified the probabilities for a given number of failures $i$ the probability of $i$ of these failures to occur must be included and then summed up over all possible $i$. With $p$, the probability that a single cell produces a failure, this is then

$$P_{1,d\geq 2}(K) = \sum_{i=0}^{\left\lfloor \frac{K}{3} \right\rfloor} p^i (1-p)^{K-i} \binom{K}{i} P_{1,d\geq 2}(K,i)$$

where $\lfloor x \rfloor$ is the floor function. For periodic boundary conditions, only the exchange $P_{1,d\geq 2}(K,i) \leftrightarrow P_{1,d\geq 2,PB}(K,i)$ needs to be made.

We have seen how to calculate the probabilities for one vertical lane of cells. To get the probability for all lanes, they must be multiplied accordingly, i.e., for $L$ vertical lanes to the $L$’th power. Additionally, it must be included that at most $L$ steps need to be taken to reach the target pattern, thus:

$$P_{d\geq 2}(L,K) = \left[ \sum_{i=0}^{\left\lfloor \frac{K}{3} \right\rfloor} p^i (1-p)^{K-i} \left( \binom{K-2i}{i} + \binom{K-2i}{i-1} + \binom{K-2i+1}{i-1} \right) \right]^{L^2}$$

$$P_{d\geq 2,\mathrm{PB}}(L,K) = \left[\sum_{i=0}^{\left\lfloor \frac{K}{3} \right\rfloor} p^i (1-p)^{K-i} \left(\binom{K-2i}{i} + 2\binom{K-2i-1}{i-1}\right)\right]^{L^2}$$

Finally, if there are no periodic boundary conditions, we can use a similar procedure to derive the probability of not having an error in the pattern (Figure S12). Additionally, the effects of the top and bottom boundaries itself must be discussed. It is assumed that the boundaries always have constant values. Averaged over all patterns and dynamics they have with $50\%$ probability the correct state and with $50\%$ probability an erroneous state. An error in the lane next to the boundary thus will manifest itself in $50\%$ of the cases. Since an error in the next lane will also be problematic after one time step (compare Fig S Figure **S12**b), this error will also manifest itself in $50\%$ of the cases. Thus, there are $4\frac{L}{2} = 2L$ cells on average in which an error will lead to an error in the target pattern. Again, it is assumed that $L$ steps need to be taken, thus the chance of not getting any error at the boundaries is

$$P_{\mathrm{Bound}} = (1-p)^{2L^2}$$

#### S4.3 Additional processes that contribute to the total error probability

The analytical approximations obtained in the previous section neglect a number of processes that affect the generation, spreading, and elimination of errors. Here, we briefly discuss these processes.

*Generation of errors directly adjacent to pre-existing errors:* In the previous section we had estimated the probability of generating two adjacent errors (which cannot be corrected) by assuming that they are generated in the same time step. However, two adjacent errors can also occur when an error survives the first correction step (as in Fig. Figure **S12**b) and an additional error is newly generated next to this pre-existing error. This process effectively increases the probability to generate two immediately neighboring errors.

*Back mutations:* There is a small chance that a point mutation reverses spontaneously in a subsequent time step. This eliminates the error and effectively decreases the overall error probability. The probability for back mutations decreases with increasing number of states *k*.

*Number of time steps:* In our estimate in the previous section, we did not take into account that different initial and target patterns have different shortest path lengths connecting them. Instead, we always used the maximum shortest path length, equal to the system size $L$, as an estimate for the actual shortest path length $t_{\mathrm{SP}}$ from a given initial pattern to a desired target pattern (which can be smaller than $L$, depending on the initial and target pattern). To include the heterogeneity in the actual shortest path lengths in the calculation of the quantity $P_{d\geq 2}(L, K)$ considered above, it would be necessary to average over the distribution of shortest path lengths.

*Correction of errors in the last steps:* As seen above, the correction of closely spaced errors may take several steps (Fig. Figure **S12**b). Therefore, the correction of such error "clusters" which appear in the last step(s) is not possible. This was not accounted for in our estimate in the previous section.

*Influence of the rules themselves:* The rules acting along the horizontal axis can influence the error correction both positively and negatively. A positive effect on error correction is obtained when the error occurs in a spatiotemporal point that cannot be reached by the steering effects of the organizers (compare previous chapter). Depending on the rule it is then shifted towards the fixed boundary and eliminated by hitting this boundary. Especially the prototypical rules 15 and 240 are designated for this behavior. In general, the properties of the rule can be read from the algebraic form. Whilst the rules 15 and 240 only shift the error and do not multiply it, the rules 90, 105 and 150 propagate the error in both directions, the latter two even leave it at the same place. Thus, rule

150 performs worst, as can be seen in the diagrams (rule 105 performs better on the boundaries, see below). Rule 60 shifts the error to the right and leaves it at the same cell. The rest of the rules shift the error to the right and, potentially (due to the nonlinearity) leave them in place and shift them left.

There can also be a large difference between the rules with regard to the boundaries, best illustrated by the rules 15 ($f = (1 + x_{i-1}^{t})\text{mod}_2$) and 240 ($f = x_{i-1}^{t}$). Whilst rule 240 only shifts the current state to the right, rule 15 also inverts the state in each step. Now consider an error appearing in the row directly below the top boundary, which is assumed to be fixed to the same state for all boundary cells. If the erroneous cell has the same state as the boundary, then the error correction mechanism cannot correct the error. For rule 240, this will then never change since the error is just shifted but has always the same state as the boundary. On the other hand, rule 15 inverts the state additional to shifting it, thus the erroneous cell will have a different state than the boundary in the next time step. Then the error correction mechanism is able to correct the error. This explains the observation that for fixed boundary conditions (Fig. SFigure **S14**), the rule 15 outperforms rule 240, but for periodic boundary conditions (Main Text, Fig. 5), there is no visible difference between the two (the data points fall perfectly on top of each other such that the ones from rule 15 cannot be seen). The properties of the rules are summed up in Table STable **S1**.

### S5 Design Concept for a Programmable DNA Cellular Automaton

One possibility of application of our model is on molecular systems such as the ones designed in the field of DNA computing. In DNA computing DNA-biopolymers are used in a smart fashion to perform computing tasks that are usually realized by an electrical circuit, often in a massively parallel way (see e.g. the work by Benenson et al.[8]).

Based on a toolkit consisting of double-stranded DNA molecules with dangling ends, ligase enzymes, and restriction enzymes, Yin et al.[9] proposed a flexible design for elementary one-dimensional cellular automata with different update rules. Here we want to propose a modification of their design that would make the DNA automata programmable. We use the nomenclature and notation of the original design and only describe the modifications.

To establish our standard programmability scheme, the organizer cell must be updated in a way that its state follows a predetermined protocol. This can be realized by additionally saving the time of the update step in the sequence of the initiator molecule. Then the first two reaction steps in the reaction wave would look like

$$I^{iT}[u] + [\bar{u}]A^{a} \rightarrow I[x]^{iTa} + [\bar{x}]^{iTa}A$$

$$I[x]^{iaT} + [\bar{x}]^{iaT}R \rightarrow I^{i'(T+1)}[u] + [\bar{u}]R$$

Additionally, there is a reaction step with an additional helper molecule needed to ensure that $A$ loses the time information:

$$H[x]^{iTa} + [\bar{x}]^{iTa}A \rightarrow H[t]^{ia} + [\bar{t}]^{ia}A$$

A change or rules can be employed by washing out the rule molecules (with all other components in solution and then flushing in a new solution with new rule molecules). Since we only use rule changes when the system is in an attractor of the dynamics, meaning there is no change in the pattern, an exact timing of the rule change is not necessary.

In summary, a DNA cellular automaton can be used to exert macroscopic control over the molecular constitution of molecules on a DNA grid after having manufactured the system. The sticky ends of the DNA molecules at the grid positions can bound to other molecules with the correct complementary end added to the solution. Therefore, such a system represents a flexible and, on a macroscale, programmable way of spatially positioning specific docking positions on a grid.

### S6 Properties of Attractor Graphs

#### S6.1 Programmability of Attractor Graphs

The first question to ask is if the patterning formation process with halting is completely programmable in the sense that it is possible to reach any pattern from any other pattern, or formulated for the attractor graph, whether this graph is strongly connected. For one-sided and embedded control, the graphs are strongly connected for grid sizes $L \in \{2,3,4,5\}$, for two-sided control this is the case for $L \in \{2,3,4,5,8\}$. Thus, the general answer for larger systems is no, the system is not completely programmable.

#### S6.2 Attractor Graphs from Outer Totalistic Rulesets

In general, outer totalistic rules acting on a certain cell are defined as rules that depend only on the current state of the cell itself and on the sum of its neighbors:

$$x_i^{t+1} = f(x_{i-1}^t, x_i^t, x_{i+1}^t) = g(x_i^t, x_{i+1}^t + x_{i-1}^t)$$

Thus, these rules are agnostic about the 'direction' from which they receive signals. Examples for such rules are rule 90 $g(x_i^t, x_{i+1}^t + x_{i-1}^t) = (x_{i-1}^t + x_{i+1}^t) mod_2$ or rule 150 $g(x_i^t, x_{i+1}^t + x_{i-1}^t) = (x_{i-1}^t + x_i^t + x_{i+1}^t) mod_2$.

When we construct attractor graphs purely from outer totalistic rulesets, calculate the number of reachable patterns and fit them with an exponential function as describe in the main text, we get for the bases of the function 1.88 (linear topology, 2 organizer cells), 1.81(linear topology, 1 organizer cells) and 1.37 (circular topology).

Whilst the values for the linear topology only slightly deviate from the values obtained using the full ruleset (1.89 and 1.82, respectively), the value for circular topology drops dramatically (from 1.89 to 1.37). This can be explained by the fact that the inherent left-right symmetry of the outer totalistic rules together with the symmetric initial state 0 prevents the left-right symmetry breaking in pattern space, no matter of the organizer input. Thus, only left-right symmetric patterns are accessible.

**S6.3 Pattern Complexity**

In the main text we have linked the complexity of a spatial pattern generated to the number of instructions (an instruction consists of a rule and associated organizer input) needed to generate it starting from a homogeneous initial condition (i.e., the configuration with all states set to 0). Here we examine some features typical of patterns at each stage in the complexity hierarchy, for simplicity only the linear topology with two organizers. At the bottom of this complexity scale will be strings requiring one instruction (Fig. S**Figure S15 | Patterns reachable for scheme which is robust against variable timing of organizer signals.** For $L = 10$ we illustrate the range in complexity displayed by patterns which can be generated by a given number of instructions under this control scheme. Clockwise from top left, we start by displaying all patterns which require only one instruction: they appear periodic in nature. Next, some of the patterns which require two and three instructions are shown, exhibiting some more interesting features. The most complicated patterns for this lattice size require seven instructions, all of which are shown. Finally, all patterns which cannot be generated for $L = 10$ are shown.Figure **S15**). There we find strings with visually simple patterns: alternating zeros and ones, domains separated by a wall. These are the strings we would intuitively expect to be most compressible. One noticeable absence however is the pattern which divides the system into two equal domains, where all states on one half of the lattice are set to zero and all states on the other half to one (i.e., configuration 31 for $L = 10$). This pattern requires two instructions to generate. In fact, the vast majority of reachable patterns can be generated with 2 to 4 instructions for $L = 10$ and as we can see in the examples these are already patterns which exhibit some nontrivial structure: domains with periodic configurations mixed with homogeneous domains (Fig. S**Figure S15 | Patterns reachable for scheme which is robust against variable timing of organizer signals.** For $L = 10$ we illustrate the range in complexity displayed by patterns which can be generated by a given number of instructions under this control scheme. Clockwise from top left, we start by displaying all patterns which require only one instruction: they appear periodic in nature. Next, some of the patterns which require two and three instructions are shown, exhibiting some more interesting features. The most complicated patterns for this lattice size require seven instructions, all of which are shown. Finally, all patterns which cannot be generated for $L = 10$ are shown.Figure **S15**). The few patterns which require a large number of instructions already exhibit an irregular spatial distribution of states and appear to be visually similar to non-reachable patterns.

### FIGURES & FIGURE LEGENDS

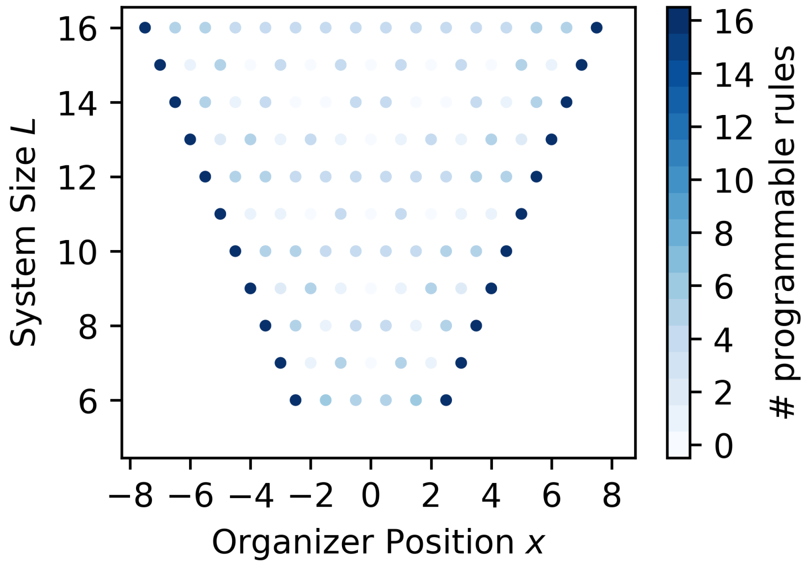

**Figure S1 | Number of programmable rules (color) as function of the position $x$ in the pattern measured from the center and the size $L$ of the system.** System sizes smaller than $L = 6$ show comparably large values and are omitted to allow for a reasonable dynamic range of the color code. The graph clearly shows that the position of the organizer cell at the boundary is optimal. Note that none of the rules remains programmable for all tested positions.

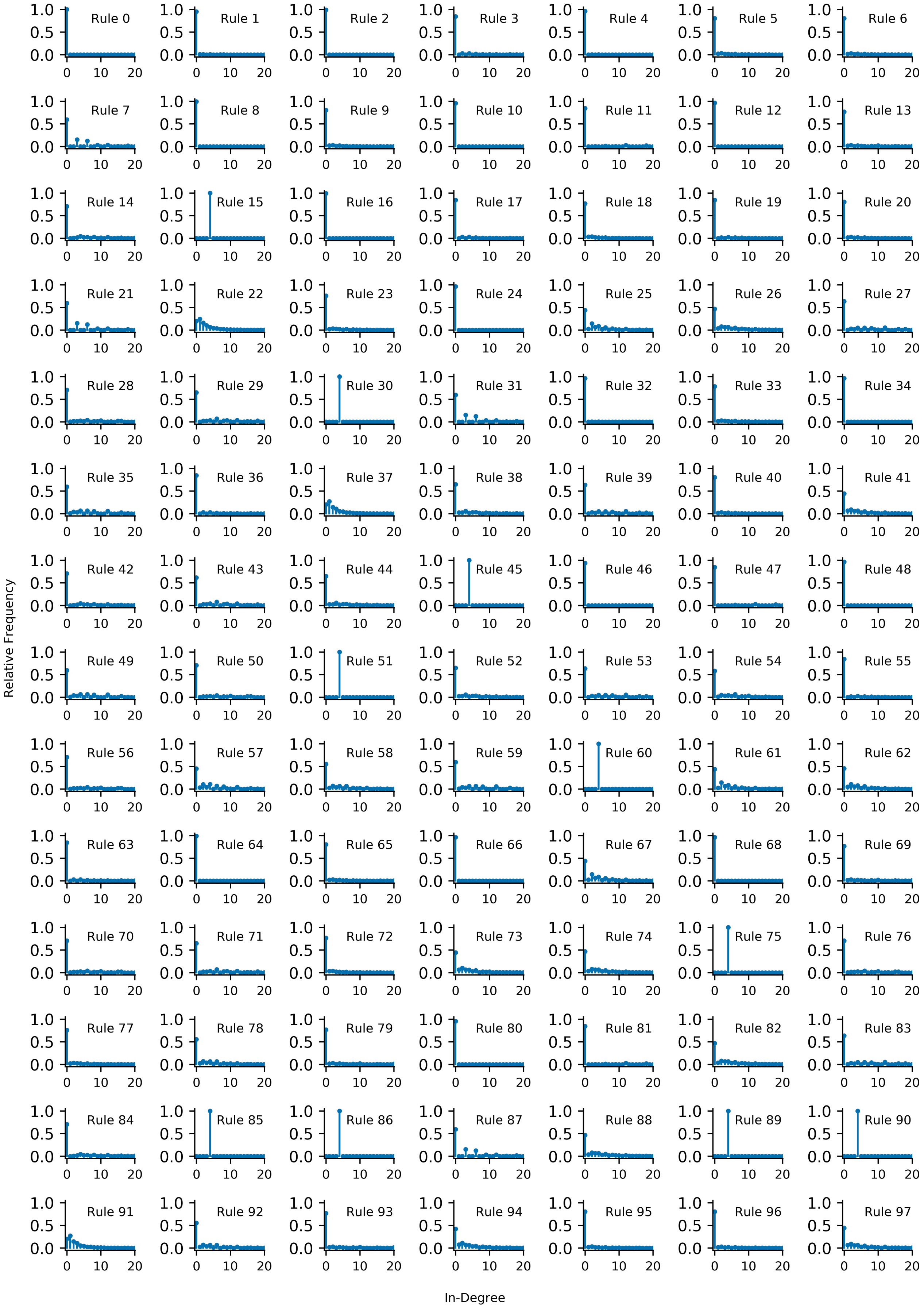
Relative Frequency
In-Degree
Rule 0
Rule 1
Rule 2
Rule 3
Rule 4
Rule 5
Rule 6
Rule 7
Rule 8
Rule 9
Rule 10
Rule 11
Rule 12
Rule 13
Rule 14
Rule 15
Rule 16
Rule 17
Rule 18
Rule 19
Rule 20
Rule 21
Rule 22
Rule 23
Rule 24
Rule 25
Rule 26
Rule 27
Rule 28
Rule 29
Rule 30
Rule 31
Rule 32
Rule 33
Rule 34
Rule 35
Rule 36
Rule 37
Rule 38
Rule 39
Rule 40
Rule 41
Rule 42
Rule 43
Rule 44
Rule 45
Rule 46
Rule 47
Rule 48
Rule 49
Rule 50
Rule 51
Rule 52
Rule 53
Rule 54
Rule 55
Rule 56
Rule 57
Rule 58
Rule 59
Rule 60
Rule 61
Rule 62
Rule 63
Rule 64
Rule 65
Rule 66
Rule 67
Rule 68
Rule 69
Rule 70
Rule 71
Rule 72
Rule 73
Rule 74
Rule 75
Rule 76
Rule 77
Rule 78
Rule 79
Rule 80
Rule 81
Rule 82
Rule 83
Rule 84
Rule 85
Rule 86
Rule 87
Rule 88
Rule 89
Rule 90
Rule 91
Rule 92
Rule 93
Rule 94
Rule 95
Rule 96
Rule 97

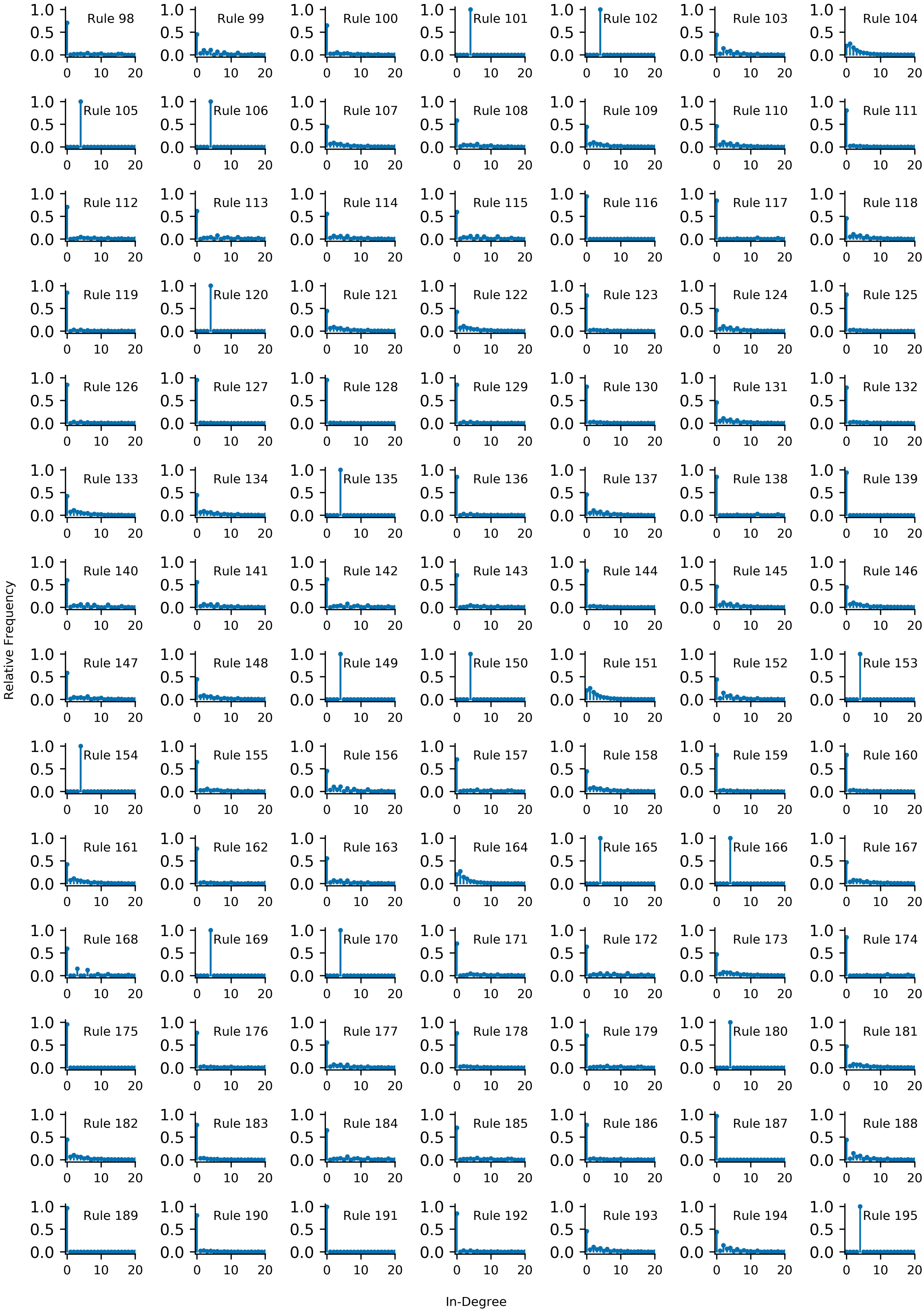
Relative Frequency
In-Degree
Rule 98
Rule 99
Rule 100
Rule 101
Rule 102
Rule 103
Rule 104
Rule 105
Rule 106
Rule 107
Rule 108
Rule 109
Rule 110
Rule 111
Rule 112
Rule 113
Rule 114
Rule 115
Rule 116
Rule 117
Rule 118
Rule 119
Rule 120
Rule 121
Rule 122
Rule 123
Rule 124
Rule 125
Rule 126
Rule 127
Rule 128
Rule 129
Rule 130
Rule 131
Rule 132
Rule 133
Rule 134
Rule 135
Rule 136
Rule 137
Rule 138
Rule 139
Rule 140
Rule 141
Rule 142
Rule 143
Rule 144
Rule 145
Rule 146
Rule 147
Rule 148
Rule 149
Rule 150
Rule 151
Rule 152
Rule 153
Rule 154
Rule 155
Rule 156
Rule 157
Rule 158
Rule 159
Rule 160
Rule 161
Rule 162
Rule 163
Rule 164
Rule 165
Rule 166
Rule 167
Rule 168
Rule 169
Rule 170
Rule 171
Rule 172
Rule 173
Rule 174
Rule 175
Rule 176
Rule 177
Rule 178
Rule 179
Rule 180
Rule 181
Rule 182
Rule 183
Rule 184
Rule 185
Rule 186
Rule 187
Rule 188
Rule 189
Rule 190
Rule 191
Rule 192
Rule 193
Rule 194
Rule 195

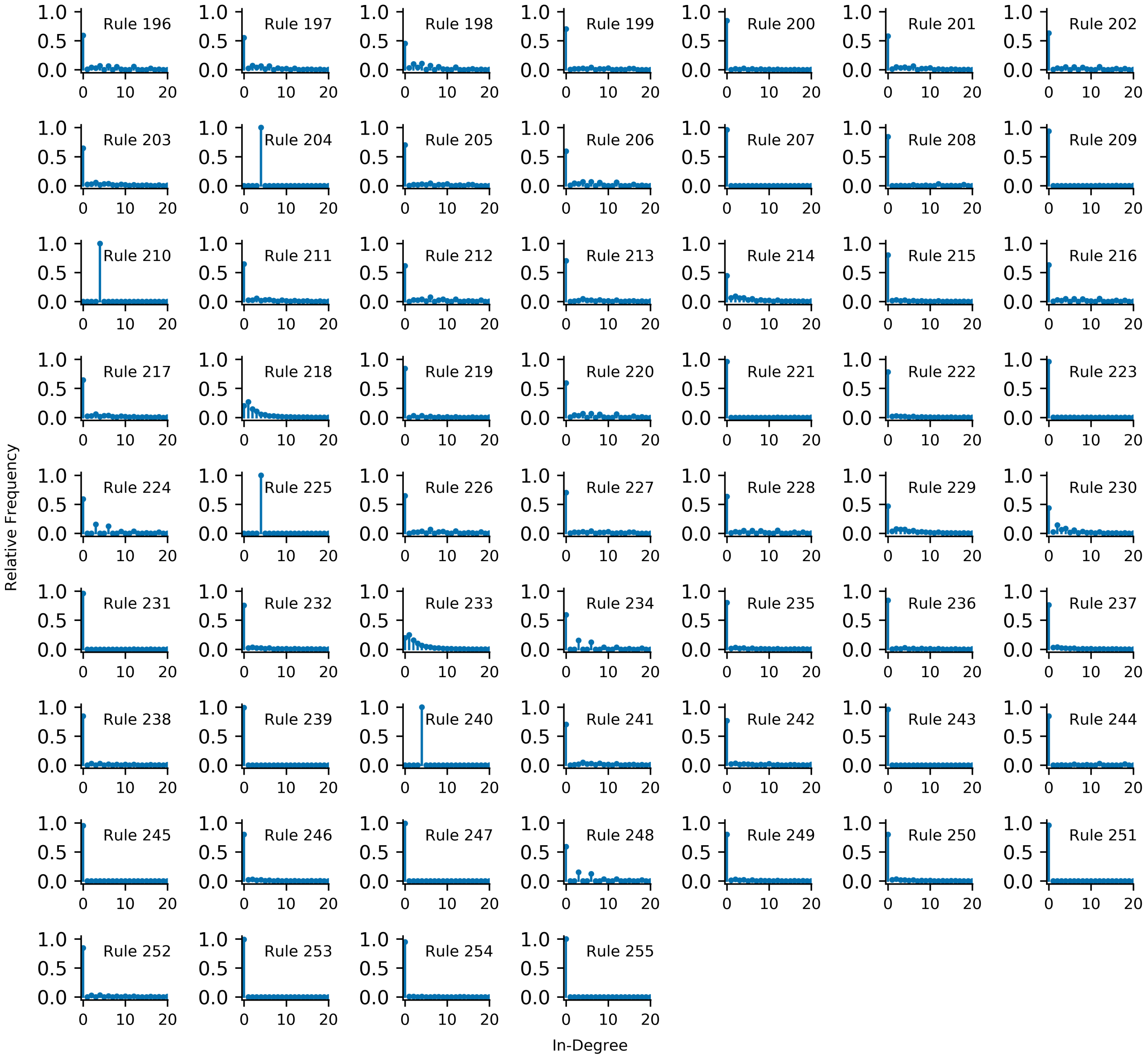

**Figure S2 | In-Degree Distributions.** In-degree frequency distribution of all rules for linear topology with 2 organizers and system size $L = 16$. The distributions are cut at an in-degree of 20. For a discussion see main text, Fig. 2f and it's caption.

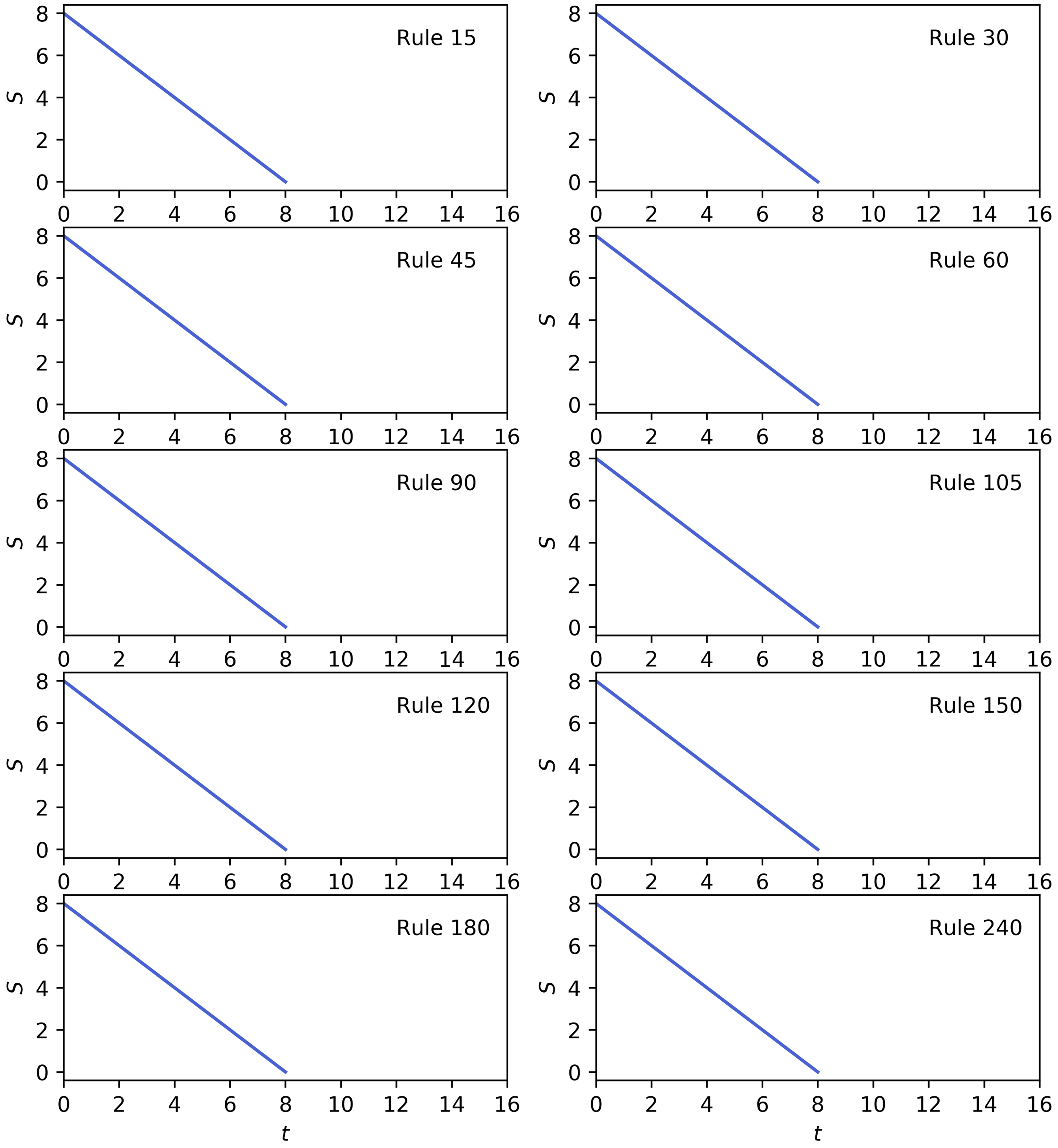

**Figure S3 | Average, minimum and maximum $S(t)$ vs. time for linear topology with an organizer cell on one side.** Plotted is the minimum, maximum, and average over all target patterns of the microcanonical entropy $S(t)$ for the programmable rules for system size $L = 8$ and linear topology with an organizer cell on one side. In this case, the minimum, maximum, and average all fall on one line.

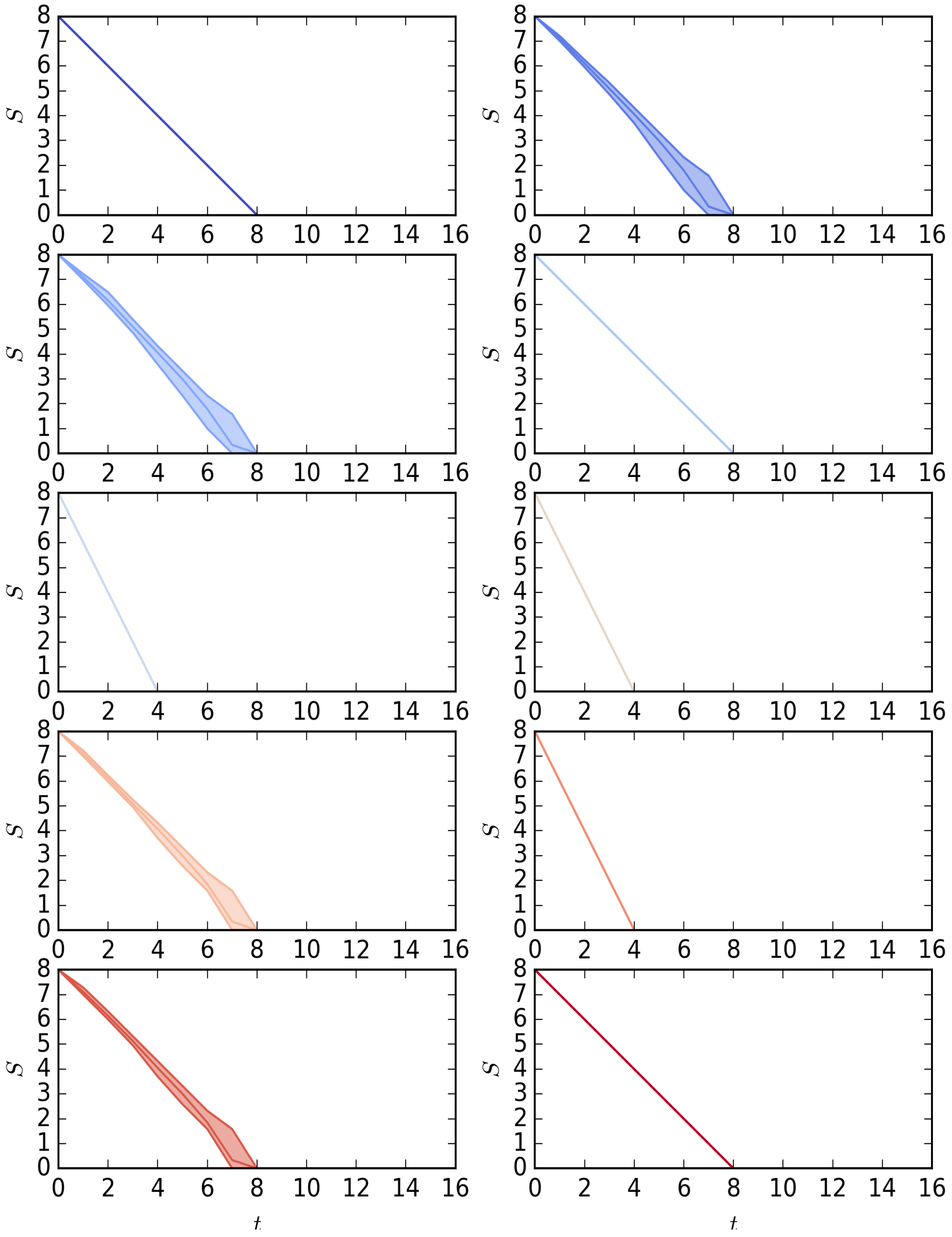

**Figure S4 | Average, minimum and maximum $S(t)$ vs. time for linear topology with organizer cells on both sides.** Plotted is the minimum, maximum, and average over all target patterns of the microcanonical entropy $S(t)$ for the programmable rules for system size $L = 8$ and linear topology with organizer cells on both sides. For the rules 15, 60, 90, 105, 150, 240, they all fall on one line.

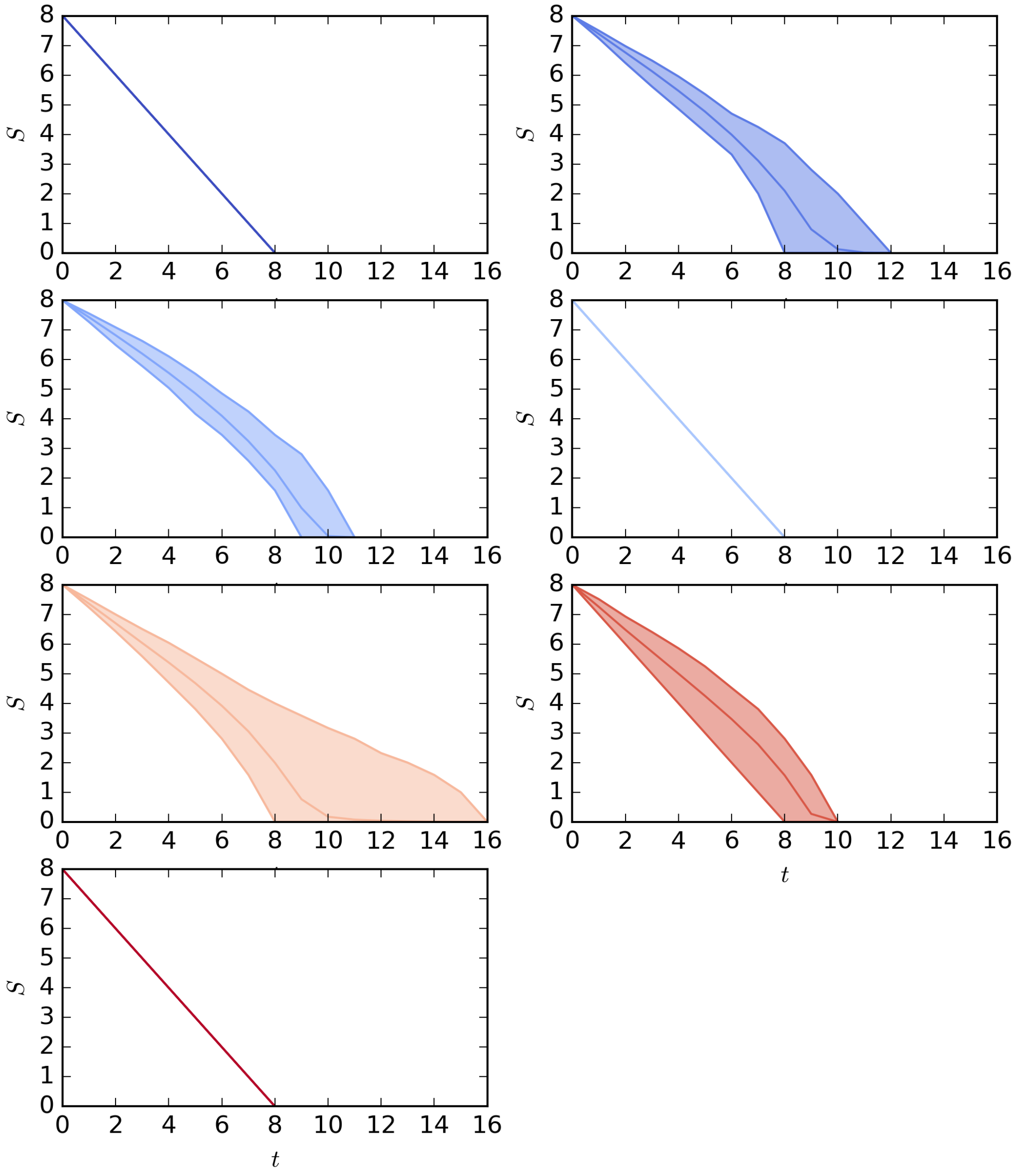

**Figure S5 | Average, minimum and maximum $S(t)$ vs. time for circular topology.** Plotted is the minimum, maximum, and average over all target patterns of the microcanonical entropy $S$ for the programmable rules, system size $L = 8$ and circular topology. For the rules 15, 60, 240, they all fall on one line.

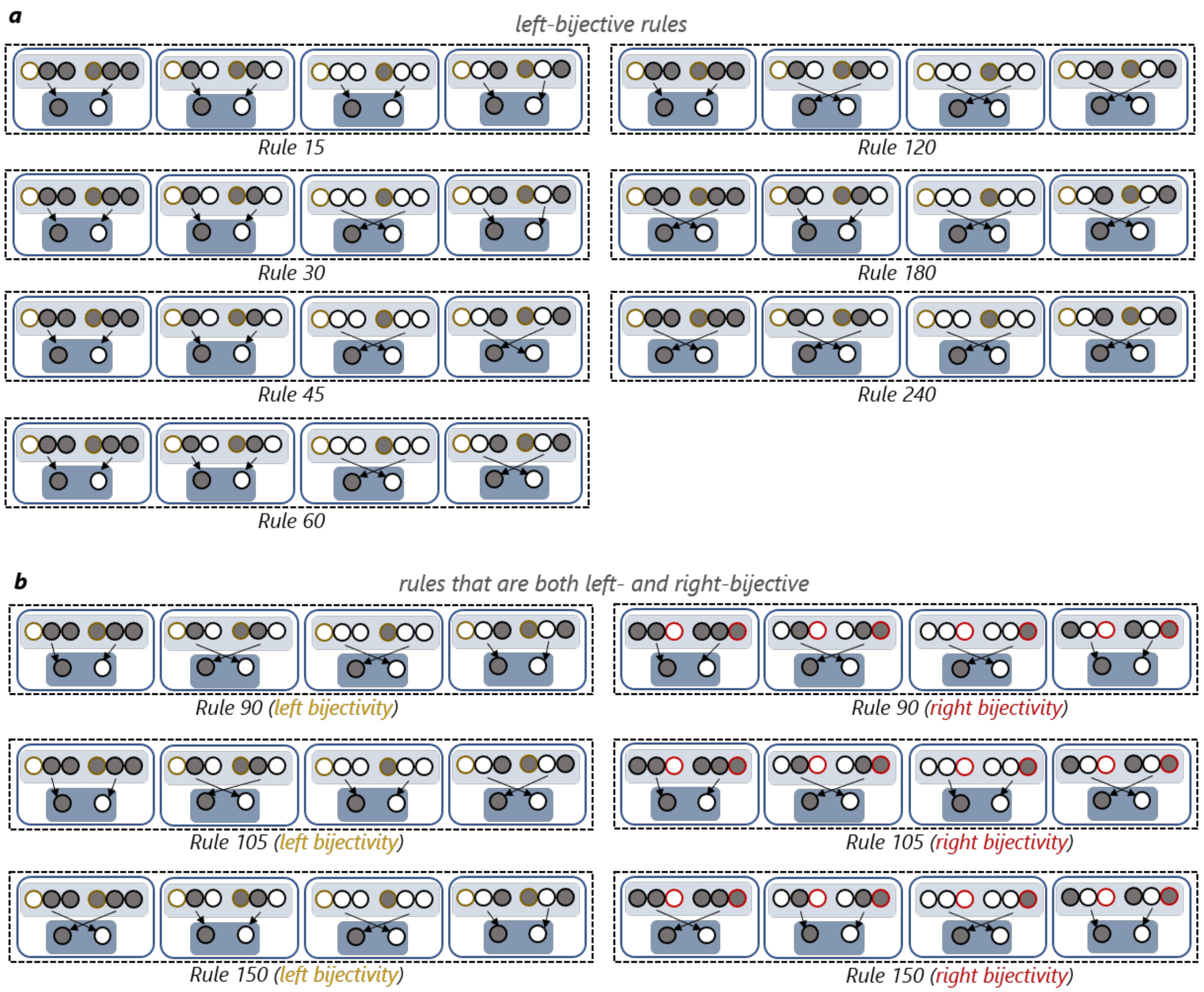

**Figure S6 | Transition tables for the programmable rules.** Design similar to Fig. 4a from the main text. The rule tables of rules $f\left(x_{i-1}(t), x_i(t), x_{i+1}(t)\right)$ which are left-bijective (a) and left- and right-bijective (b) are grouped according to their middle and right inputs $x_i(t), x_{i+1}(t)$ ((a) and left part of (b)) or according to their left and middle inputs $x_{i-1}(t), x_i(t)$ (right part of (b)) and demonstrate that if the remaining input changes, the output changes, thus demonstrating bijectiveness.

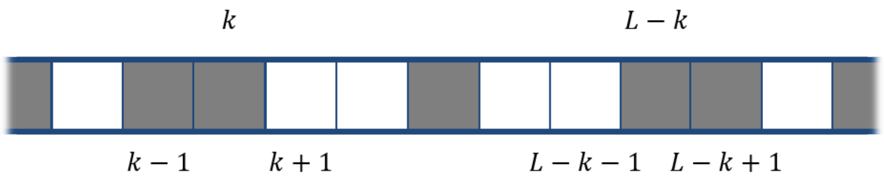

**Figure S7 | Illustration of the symmetry argument.** Shown is a part of the symmetric pattern with the labels used in the text.

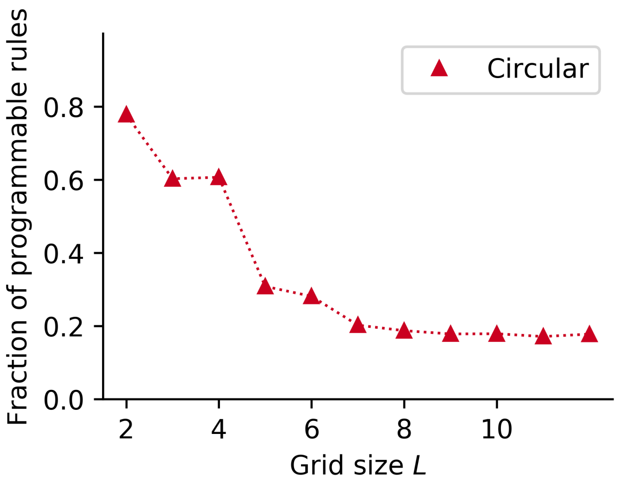

**Figure S8 | Fraction of Programmable Rules for Circular Topology**. Depicted is the fraction of the programmable rules out of a randomly chosen subset of the rules $f(x_{i-1}, x_i, x_{i+1})$ which are bijective in one of their outer arguments $x_{i-1}$, $x_{i+1}$ for a number of states $k = 3$. Fraction for the largest tested grid size $L = 12$ is 17.8%

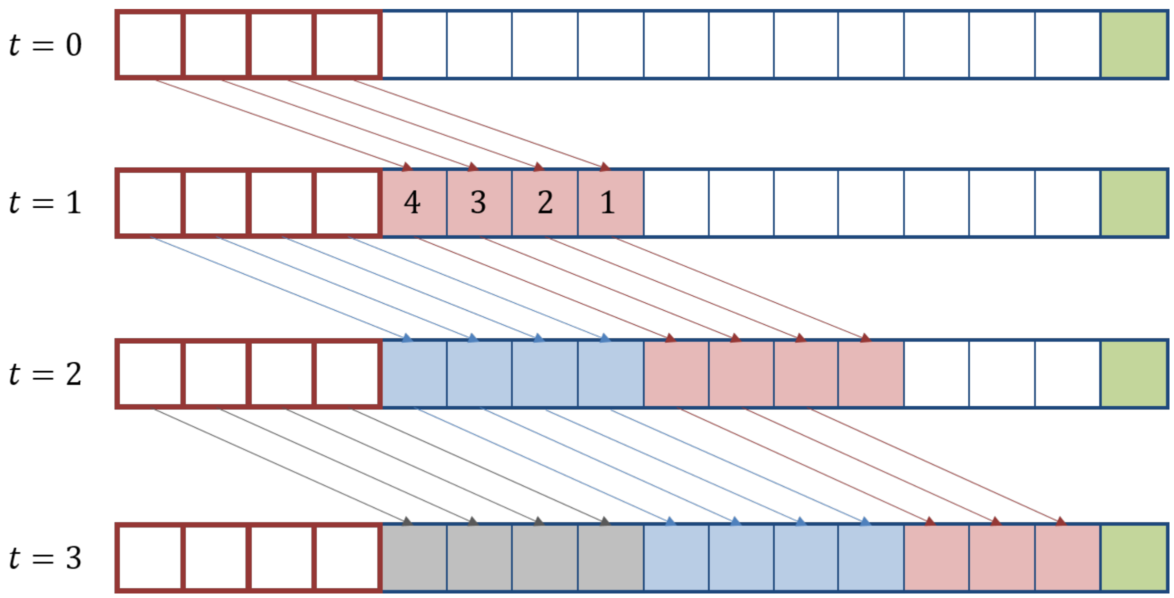

**Figure S9 | Illustration of programmability in cellular automata with larger neighborhoods.** Shown is a schematic illustration of a cellular automaton with a function $f = f(x_{i-4}, x_{i-3}, x_{i-2,}, x_{i-1,} x_i, x_{i+1})$ which is bijective in the arguments on the left side. Since the radius of the rule is then $r = 4$, $4$ organizer cells indicated through red boxes are needed. The arrows indicate which cell is used to influence which cell at a later time $t$. The numbers in the cells at $t = 1$ show exemplary what the order is in which the cells need to be set when the values of the organizer cells are determined. First, the innermost cell needs to be set because it is the only one for which the whole neighborhood is already known. For setting cell $2$ first, for example, the value of the organizer cell at neighborhood position $x_{i-3}$ is not known. The rest of the logic follows the previously demonstrated proof. Further, it is illustrated that depending on the grid size and the radius of the rule in the last step the full potential of determining $r$ cells at each step is not needed and the number of steps is given by $\left\lceil \frac{L}{r} \right\rceil = \left\lceil \frac{11}{4} \right\rceil = 3$ here.

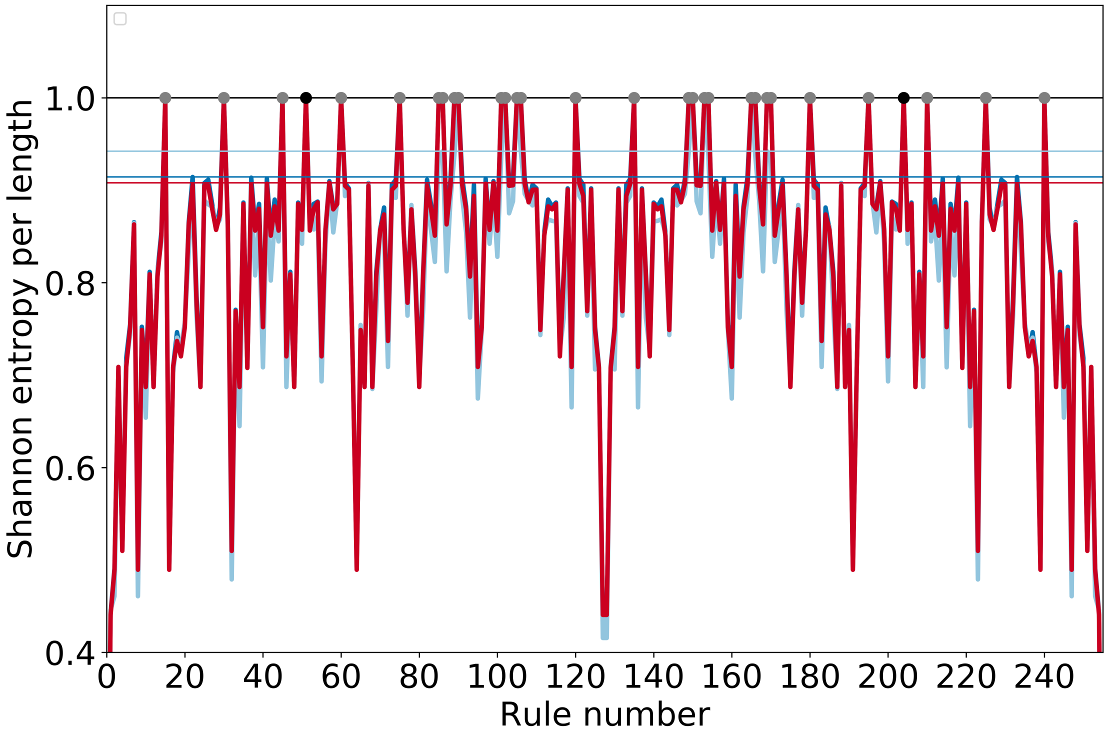

**Figure S10: Shannon Entropy per Length (16) for all elementary cellular automata rules and topologies.** The Shannon entropy per length $\frac{H}{L} = -\frac{1}{L}\sum_i p_i\, log_2(p_i)$ with $p_i = \frac{\text{InDegree(Node } i)}{\sum_j \text{InDegree(Node } j)}$. The different topologies are indicated by different colored thick lines (dark blue – linear, 2 organizers; light blue – linear, 1 organizer; red - circular). Please note that the lines are just drawn to make the figure more readable but shall not indicate some continuous connection between to neighboring datapoints. The thin lines in the corresponding color indicate for each case the largest value smaller than $\frac{H}{L} = 1$. The grey-colored dots show mark the programmable rules, the two black colored dots the two non-programmable rules scoring $\frac{H}{L} = 1$. It can be seen that nearly all the rules that score a value $\frac{H}{L} = 1$ belong to the class of programmable rules. The two exceptions can trivially be identified as being not programmable since they are the *identity* (204) and the *complement* (51) rule.

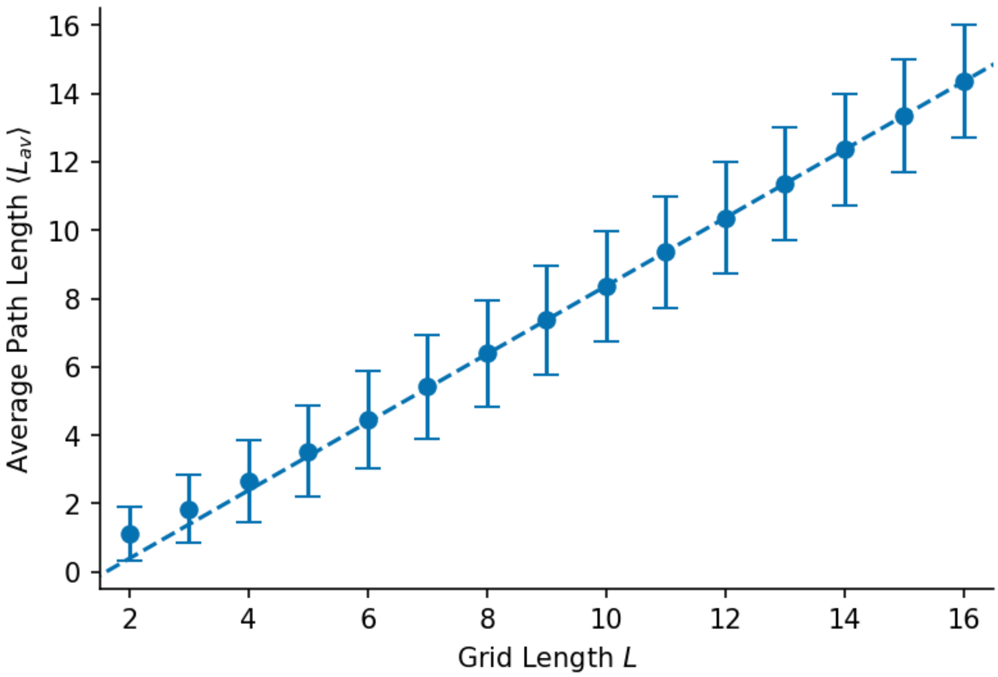

**Figure S11 | Fit of the average path length.** Plotted is the average path length for the programmable rule 45 and linear topology with one organizer cell as a function of the grid length, together with the linear fit function.

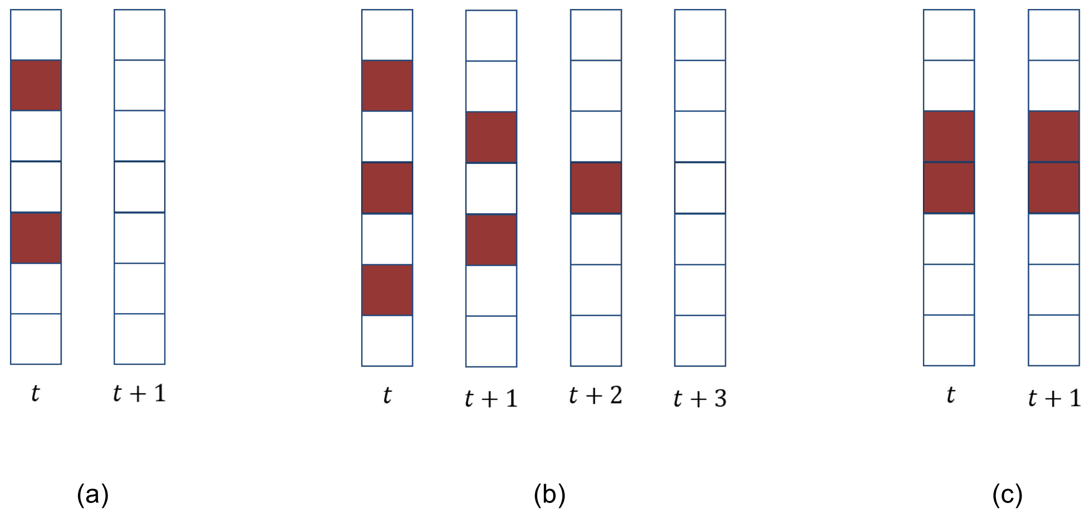

**Figure S12 | Illustration of the error correction dynamics |** Shown are different time evolutions of errors (red cells). (a) If two errors are well separated, they are both corrected in a single step by the majority voting rule. (b) Even if multiple errors are close, but separated by at least one cell, they are still corrected after a few time steps. (c) If two errors are directly adjacent, they will persist and cannot be corrected by the majority voting rule.

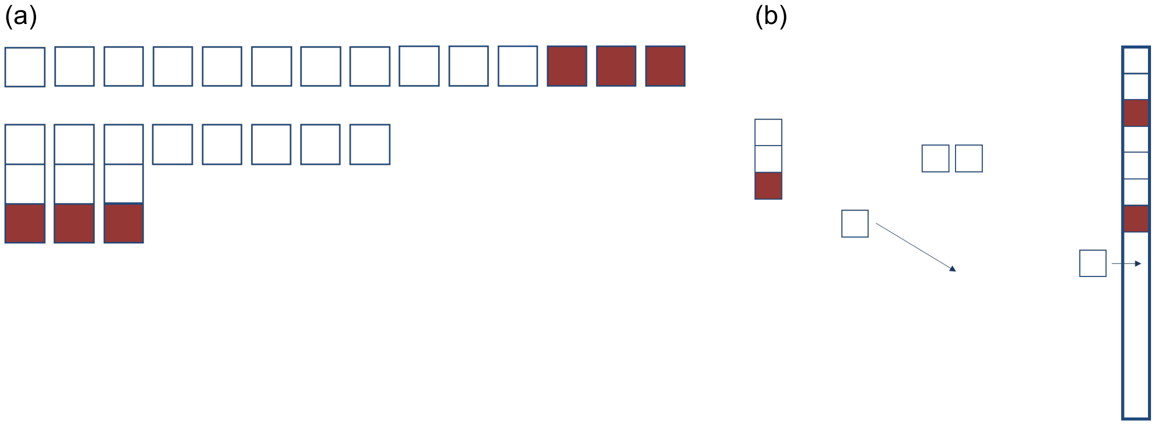

**Figure S13 | Illustration of combinatorial calculations |** (a) Displayed in the first row are the $K - i$ correct cells (white) and $i$ errors (red). They will be combined to $i$ elements of one error and one correct cell, leaving $K - 2i$ non-errors (second row). These will be combined into a vertical line (b), ensuring that no two errors touch each other directly.

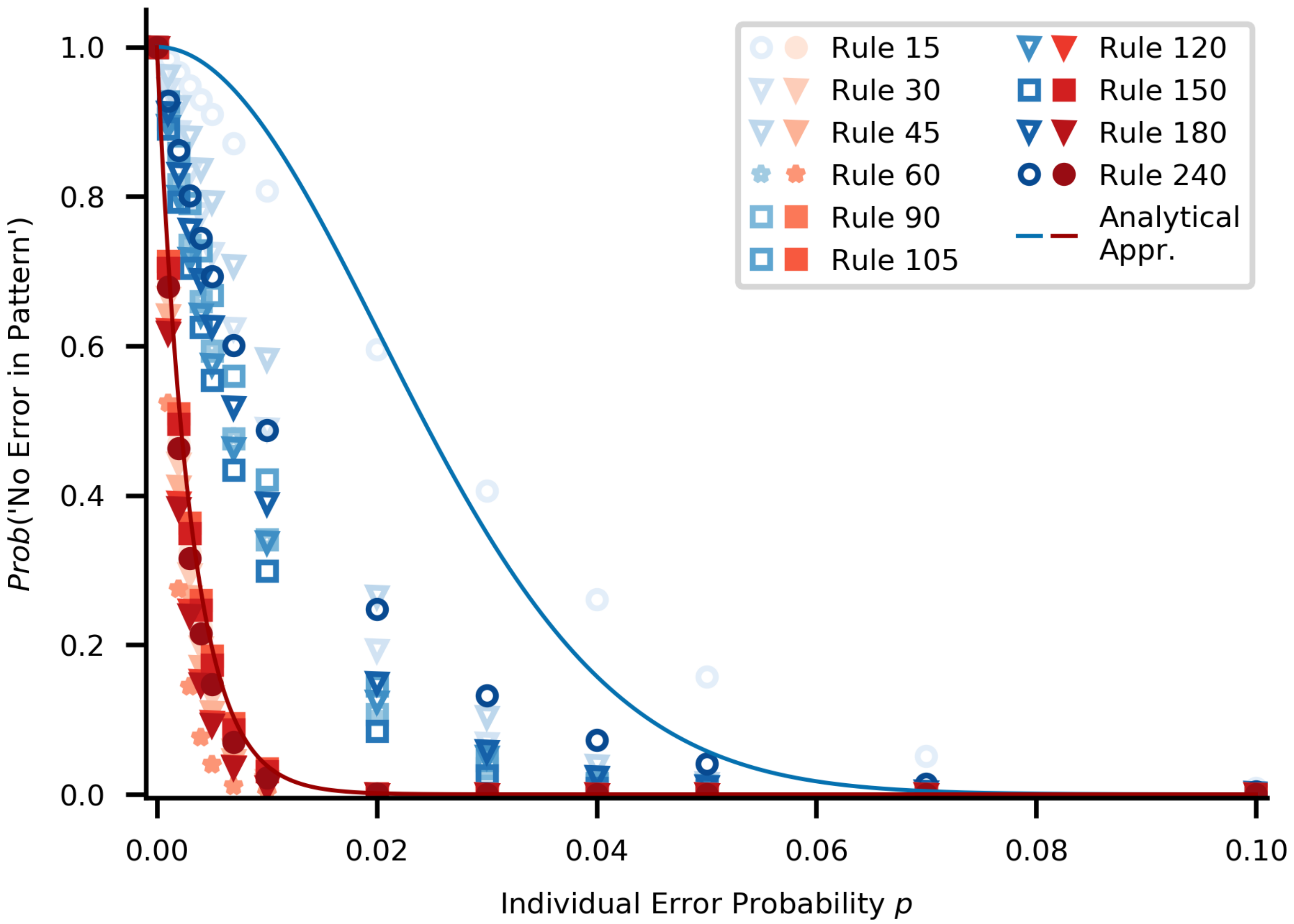

**Figure S14 | Error Robustness in a planar system.** Same plot as Fig. 5c in the main text, but for a planar grid of cells instead of the tubular topology. In the planar system, the boundary states are homogeneous and constant in time.

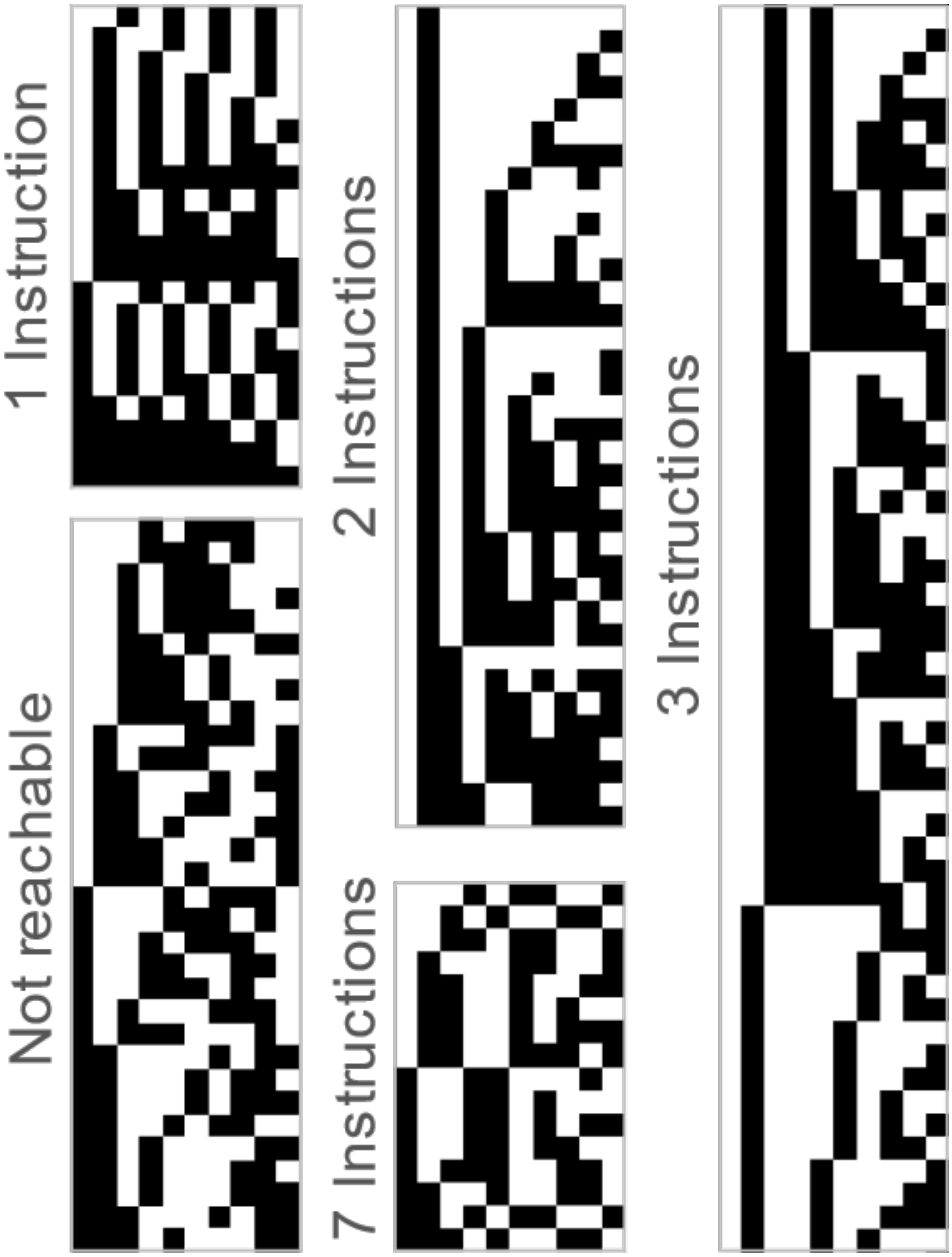

**Figure S15 | Patterns reachable for scheme which is robust against variable timing of organizer signals.** For $L = 10$ we illustrate the range in complexity displayed by patterns which can be generated by a given number of instructions under this control scheme. Clockwise from top left, we start by displaying all patterns which require only one instruction: they appear periodic in nature. Next, some of the patterns which require two and three instructions are shown, exhibiting some more interesting features. The most complicated patterns for this lattice size require seven instructions, all of which are shown. Finally, all patterns which cannot be generated for $L = 10$ are shown.

### SUPPLEMENTARY TABLES

**Table S1 | Error propagation properties of the programmable rules for linear topology with one organizer cell on the left.** For all programmable rules, the algebraic form and the error propagation properties derived from it are displayed.

| Rule | Algebraic Form $f(x_{i-1}^t, x_i^t, x_{i+1}^t)$ | Error propagation | | |
|---|---|---|---|---|
| | | Shifted left | Stays | Shifted right |
| **15** | $(1 + x_{i-1}^t)\ \mathrm{mod}_2$ | No | No | Yes |
| **30** | $(x_{i-1}^t + x_i^t + x_{i+1}^t + x_i^t x_{i+1}^t)\mathrm{mod}_2$ | Potentially | Potentially | Yes |
| **45** | $(1 + x_{i-1}^t + r + x_i^t x_{i+1}^t)\mathrm{mod}_2$ | Potentially | Potentially | Yes |
| **60** | $(x_{i-1}^t + x_i^t)\mathrm{mod}_2$ | No | Yes | Yes |
| **90** | $(x_{i-1}^t + x_{i+1}^t)\mathrm{mod}_2$ | Yes | No | Yes |
| **105** | $(1 + x_{i-1}^t + x_i^t + x_{i+1}^t)\mathrm{mod}_2$ | Yes | Yes | Yes |
| **120** | $(x_{i-1}^t + x_i^t x_{i+1}^t)\mathrm{mod}_2$ | Potentially | Potentially | Yes |
| **150** | $(x_{i-1}^t + x_i^t + x_{i+1}^t\ )\mathrm{mod}_2$ | Yes | Yes | Yes |
| **180** | $(x_{i-1}^t + x_i^t + x_i^t x_{i+1}^t)\mathrm{mod}_2$ | Potentially | Potentially | Yes |
| **240** | $x_{i-1}^t$ | No | No | Yes |
